# A cell surface code mediates tissue-intrinsic defense against aberrant cells in epithelia

**DOI:** 10.1101/2023.02.16.528665

**Authors:** Friedericke Fischer, Laurin Ernst, Anna Frey, Katrin Holstein, Deepti Prasad, Vanessa Weichselberger, Ramya Balaji, Anne-Kathrin Classen

## Abstract

Tissue-intrinsic error-correction mechanisms allow epithelial cells to detect aberrant neighboring cells and cause their removal from the tissue. The molecular mechanisms which grant cells the ability to compare their internal states is unknown. Here we demonstrate that comparison of cell identity, created by cell-fate-specifying transcription factors and patterning pathways, is conveyed through a specific set of cell surface molecules. We demonstrate that Drosophila imaginal discs express a range of cell surface molecules previously implicated in neuronal axon guidance processes, such as members of the Robo, Teneurin, Ephrin, Toll-like or atypical Cadherin families. Expression of these molecules is regulated by intrinsic fate-patterning pathways of the disc but also by aberrant expression of oncogenic Ras^V12^. Importantly, mosaic clones deregulating individual cell surface molecules are sufficient to induce all hallmarks of ’interface surveillance’, a tissue-intrinsic error-correction mechanism previously shown to be induced by cells with aberrant activation of fate-patterning pathways. Specifically, cells with deregulated expression of Robo2 and Robo3 induce actomyosin enrichment, bilateral JNK signaling and apoptosis at mosaic clone interfaces in imaginal discs. Moreover, deregulation of Robo2 levels, which is normally expressed in a complex endogenous pattern, induces these interface surveillance hallmarks in a Robo2-pattern-specific manner. Taken together, our work indicates that these cell surface molecules mediate cell fate recognition in epithelial tissues and thereby contribute to the maintenance of epithelial health by initiating detection and removal of aberrant cells during development and adult tissue homeostasis.

## Introduction

Genetically altered cells repeatedly appear in epithelial tissues, either because of developmental errors or mutagenesis ^1, 2^. Surveillance and removal of these cells is required to maintain tissue homeostasis and organismal health. ’Interface surveillance’ - a distinct branch of tissue-intrinsic error-correction mechanisms like cell competition - mediates removal of aberrant cells from epithelial tissues ^3, 4^. ’Interface surveillance’ responses are specifically driven by aberrant cells that carry mutations in cellular signaling pathways or transcriptional networks which control cell fate and differentiation programs. For example, mutations in patterning pathways (Dpp/TGF-1, Wg/WNT, Hh/Shh, JAK/STAT, Notch) or in cell-fate-specifying transcription factors (Arm, Iro-C, Omb, Yki, En/Inv, Ap, Ci, Hox genes) induce phenotypes related to ’interface surveillance’ ^5–23^. Mosaic clones deregulating these pathways create a pronounced difference in cell fate between ’aberrant’ and surrounding ’normal’ cells, and we previously demonstrated that such steep differences in fate between neighboring cells causes enrichment of Myosin II and filamentous Actin at shared junctional and lateral interfaces in Drosophila imaginal epithelia. This response drives cell segregation between two cell populations via smoothening of the contractile interface, a characteristic hallmark of ’interface surveillance’ ^3^. Importantly, this response is induced in a strict position-dependent manner according to the cell fate of the surrounding cells. For example, Cubitus interruptus (Ci)-expressing clones have normal corrugated shapes in anterior wing compartments, where Ci-activation by Hedgehog (Hh) signaling is high. However, Ci-expressing clones undergo clone smoothening and die in posterior compartments, where Ci-signaling is normally low ^3^. Curiously, actomyosin recruitment and interface smoothening bears a striking resemblance to compartment boundary formation in developing tissues, where two cell populations of distinct cell fate mechanically segregate via the formation of a contractile actomyosin interface between them ^24–26^. However, a distinct feature of ’interface surveillance’ - when ’normal’ cells are juxtaposed to ’aberrant’ cells - is that actomyosin recruitment is also accompanied by elevated apoptosis. Apoptosis is induced by activation of pro-apoptotic JNK signaling in cells on either side of the clonal interface ^3, 4^. This drives elimination of aberrant cells and is thus an essential hallmark of ’interface surveillance’ and its ability to maintain tissue health.

Despite this work, it is not yet understood how cells recognize that a neighbor acquires a very distinct and thus potentially aberrant fate. Recruitment of actomyosin to the contact interface and bilateral JNK activation in cells on both sides of the interface indicates that surveillance of neighbors is mediated by a cell-contact and thus cell-surface-dependent mechanism. The molecular machinery acting in interface surveillance must therefore be composed of molecules, (1) which can modulate actomyosin function and (2) which are competent to transduce contact-dependent signals into cellular signaling pathways, such as JNK. In addition, (3) expression of these molecules must be regulated by cell-fate specifying transcription factors and pathways, thereby encoding cell identity. Classical cadherins, mediating cell adhesion at adherens junctions, are prime examples of transmembrane molecules that may fulfill these criteria ^27^. Yet, the striking pattern-specific activation of ’interface surveillance’ for multiple fate-patterning pathways in imaginal discs, as described for Ci-expressing clones above, strongly implicated that more than one molecule is necessary to distinguish any two of many possible cell fates from each other ^3, 4^. A class of well-studied molecules have recently emerged to fulfil novel and unexpected roles in epithelial tissues. Specifically, members of the neuronal axon guidance and neuronal adhesion families are receiving increasing attention for their physiological and pathological roles in epithelial tissues ^28, 29^. Axon guidance molecules generally mediate signaling via interaction of a transmembrane receptor with a cognate ligand. Specifically, Roundabout (Robo), Plexin, Frazzled (Fra) or Ephrin (Eph) receptor families interact with respective ligands of the Slit, Semaphorin (Sema), Netrin (Net) and Ephrin families ^30–33^. More recent work suggests that Teneurins interact with Leucine-rich repeat (LRR)-domain proteins on neighboring cell surfaces ^33, 34^. Extensive research on axon guidance effectors has revealed this general principle: binding of the ligand to the receptor elicits signaling to recruit actomyosin modulators of the small GTPase family (Rho, Rac) and members of the actin-modulating Ena/Wasp, Abl or Src pathways ^32^. This leads to either formation or retraction of actin-based cellular protrusions, thereby driving attractive or repulsive behaviors during axon pathfinding and mediating appropriate spatial positioning of cells within neuronal systems. It is conceivable, that expression and modulation of these molecules may also have a strong impact on actomyosin and adhesion dynamics in an epithelial tissue thereby regulating epithelial morphogenesis and homeostasis ^29, 30, 35^. Indeed, Ephrin, Teneurins and LRR-domain proteins including Toll-like receptors have been strongly implicated in mechanical separation of cell populations at compartment boundaries via regulation of a contractile actomyosin interface in epithelial tissues of vertebrate and invertebrate species ^24–26, 36, 37^.

The phenomenological similarities lead us to explore if these molecules played a role in interface surveillance against aberrant cells. Importantly, the diverse fates created by patterning pathways in the imaginal disc made this tissue an ideal system to experimentally address the cell recognition system depending on fate specification. We find that neuronal cell surface molecules are widely expressed in epithelia, that a single surface molecule is sufficient to induces all hallmarks of interface surveillance responses and that these molecules are regulated by cell fates.

## Results

### Cell surface molecules with described roles in axon guidance are expressed in wing disc epithelia

To begin to understand if epithelial tissues may use cell surface molecules to distinguish fates and detect aberrant cells, we first asked if proteins functioning in axon guidance, neuronal targeting and cell adhesion are expressed in wing imaginal discs. We analyzed expression of these cell surface molecules in early and late wing disc development using published tools, such as antibodies, enhancer traps or GFP-tagged proteins ^38, 39^. To provide a reference for spatial position and developmental compartments, we included a staining for Patched (Ptc, anterior-posterior boundary) and Wingless (Wg, dorsal-ventral boundary). We were surprised to find that the majority of examined molecules were indeed expressed in wing discs, with increasingly complex expression patterns towards late 3^rd^ instar stages (**Fig 1 A-E**, **Fig S1A**). Moreover, certain attributes of those patterns, such as an elevated expression in the central pouch or the hinge, indicate regulation by central wing disc specification pathways activated by Wingless (Wg), Hedgehog (Hh), Decapentaplegic (Dpp) or Unpaired 1 (Upd1) (**Fig S1B**) ^40^. Consequently, each spatial position within the disc is endowed with a unique cell surface molecule expression profile (**Fig S1C**). The wide-spread expression of neuronal cell surface molecules suggests that these proteins may be linked to uncharacterized functions during development of imaginal discs.

**Figure 1.**
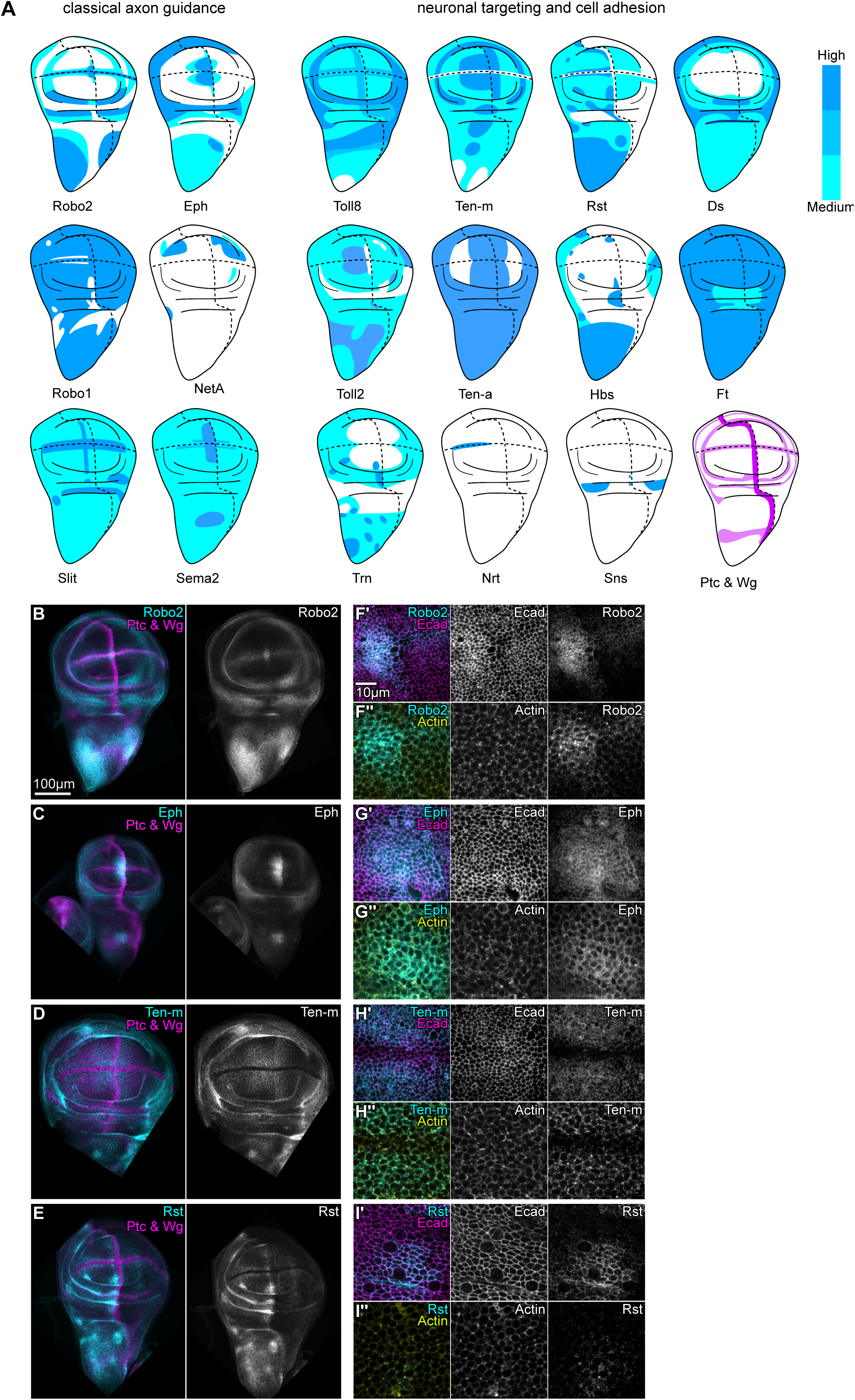
Cell surface molecules with described roles in axon guidance are expressed in wing disc epithelia. **A** Illustrations of expression patterns of cell surface molecules in third instar wing discs at 102 h AEL. Relative fluorescence intensity is depicted by color intensity from low (white), medium (cyan) to high (blue), see Fig S1 and Experimental procedures. Expression domains of compartment boundary markers Patched (Ptc, expressed in anterior cells at the anterior/posterior compartment boundary) and Wingless (Wg, expressed in cells on both sides of the dorsal/ventral compartment boundary) are shown in magenta and light magenta, respectively. All illustrated expression patterns were observed in n ≥ 4 wing discs in n ≥ 2 experimental replicates. **B-E** Wing discs expressing fusion proteins from *robo2-GFP*, *eph-GFP, ten-m-GFP* and *rst-GFP* constructs under native regulatory control (grey or magenta). Ptc and Wg demarcate A/P and D/V compartment boundaries. Images are shown at same scale. **F’-I’’** Pouch domains of *robo2-GFP*, *eph-GFP, ten-m-GFP*, and *rst-GFP* expressing wing discs (grey or cyan), also stained for the adherens junction marker E-cadherin (Ecad, grey or magenta) and cortical F-Actin (by Phalloidin, grey or yellow). Apical (’) and lateral (’’) local-z-projections, displaying receptor localization relative to junctional Ecad and relative to more lateral domains. Images are shown at same scale.

To confirm that these proteins indeed locate to the cell surface in wing disc epithelia, we analyzed their subcellular distribution in more detail. We found that Roundabout 2 (Robo2), Ephrin receptor (Eph), Teneurin-m (Ten-m) or Roughest (Rst) localized to adherens junctions and lateral domains of wing epithelial cells (**Fig 1 F’-I’’**). Both adherens junctions and lateral surfaces establish direct contact between neighboring epithelial cells, and it is these cellular surfaces that were previously shown to respond to the presence of neighboring aberrant cells, as they specifically recruit actomyosin. Thus, Robo2, Eph, Ten-m and Rst localize to cell surfaces that are implicated in activation of interface surveillance ^3, 4^. Combined, these observations open the possibility that epithelia may utilize combinatorial expression of cell surface molecules localizing to adherens junctions and lateral surfaces to detect the fate and identity of neighboring cells.

### Cell surface molecules with described roles in axon guidance mediate cell-cell recognition in wing discs

To understand if these cell surface molecules indeed play a role in aberrant cell recognition, we asked if misexpression of any of them in a mosaic context is sufficient to induce interface surveillance hallmarks. Mosaic clones expressing aberrant fates generally induce actomyosin recruitment, apoptosis, and importantly bilateral JNK interface signaling (**Fig S2.1 A-E**) ^4^. We thus used bilateral activation of the JNK-activity reporter TRE-RFP as a read-out for a targeted genetic screen ^41^. Where possible, we analyzed overexpression, as well as RNAi-mediated downregulation of cell surface molecules using the mosaic GAL4/UAS flp-out system ^42^. Strikingly, we found that deregulation of individual members of 6 cell surface protein families was each sufficient to induce TRE-RFP activation at clone boundaries in wing imaginal discs. Specifically, ectopic up-and downregulation of members of the Robo family (Robo 2 and 3), the Teneurin family (Ten-a, Ten-m), atypical Cadherins (Fat, Ds), the Eph/Ephrin system, Netrins (Netrin-B), and the LRR-proteins Toll-2, Toll-8 and Tartan was sufficient to induce bilateral JNK interface signaling (**Fig 2**). Importantly, several protein families, such as members of the Plexin/Semaphorin, IRM and Dscams families failed to score positively for JNK interface signaling in our screen, suggesting that not all cell surface molecules with roles in neuronal processes are involved in interface surveillance responses (**Fig S2.1F**, **Fig S2.2**). We conclude that a distinct set of cell surface molecules can mediate bilateral JNK signaling, otherwise a characteristic hallmark of interface surveillance of cells with aberrant fate and specification programs. We thus wanted to examine if these cell surface molecules may act as cell fate sensors required for the induction of the interface surveillance response.

**Figure 2.**
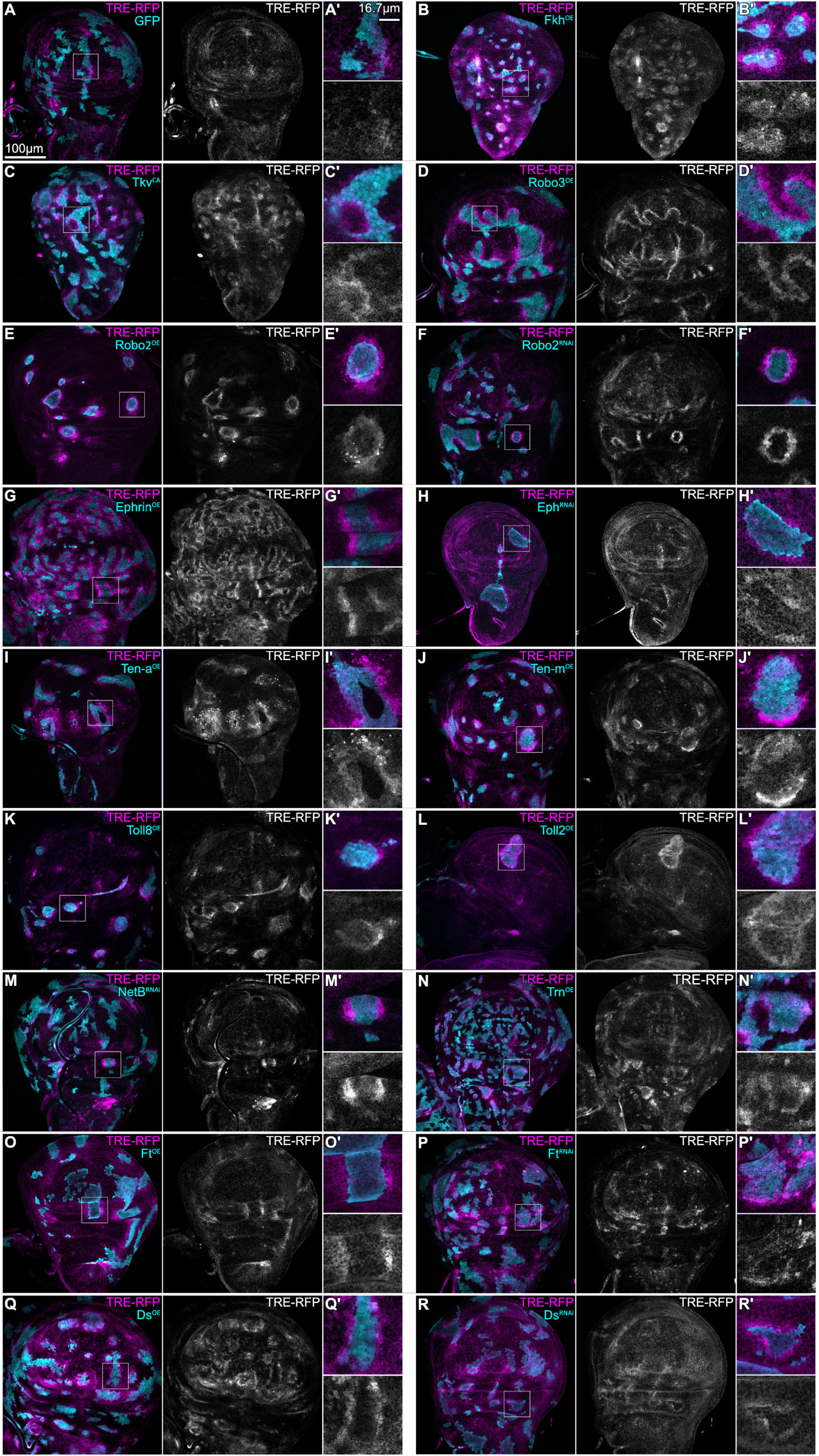
Epithelial cell surface molecules with described roles in axon guidance mediate cell-cell recognition in wing discs. TRE-RFP expression is reporting JNK pathway activity (grey or magenta in all panels). **A** A mosaic wing disc with wild type clones expressing *UAS-GFP* (GFP, cyan). Please note that JNK activity is physiologically elevated at the A/P compartment boundary in wing discs at this developmental stage. White frame marks region shown in (A’). **B, C** Mosaic wing discs with clonal alterations in fate specifying pathways by overexpression of Fkh (Fkh^OE^) and expression of constitutively active Tkv (Tkv^CA^) using *UAS-*constructs. Please note induction of bilateral JNK-signaling at mosaic interfaces. White frame marks regions shown in (B’,C’). **D-R** Mosaic wing discs with aberrant expression of epithelial cell surface molecules. Clones express GFP (cyan) and were exposed to overexpression (^OE^) by *UAS-*constructs or downregulation by *UAS-RNAi* (^RNAi^) constructs of epithelial cell surface molecules, as indicated in all panels. White frame marks regions shown in (D’-R’). Images in A-R are shown at the same scale. JNK responses in A-C have been observed in n ≥ 3 experimental replicates for this study and previous studies ^4^. JNK responses in F, H and J-L have been observed in n ≥ 3 experimental replicates for this study. JNK response in G, I and M-R has been observed in n ≥ 3 wing discs in one experimental replicate, which corresponds to the initial candidate screen (see Fig S2.1).

### A mismatch of Robo2 or Robo3 receptor expression levels between neighboring cells is detected by interface surveillance in wing discs

To analyze the results from our screen more closely, we focused on the role of Robo2 and Robo3. While atypical Cadherins, Teneurins, Ephrin and LRR-proteins that were analyzed in our screen have emerging or even established roles in epithelial tissues, almost nothing is known about the epithelial function of Robo receptors ^36, 37, 43–45^. This led us to pursue their analysis. We thus first analyzed if deregulation of Robo3 induces all hallmarks of interface surveillance, including actomyosin enrichment and apoptosis in interface cells. Robo3 is expressed at barely detectable levels in the wing imaginal disc (**Fig S3A**). Hence, overexpression of Robo3 should strongly induce interface surveillance, as Robo3-overexpressing clones would create pronounced differences in Robo3 levels to surrounding wild type cells. Indeed, Robo3-expressing clones induced JNK interface signaling, Actin enrichment at interfaces, clone smoothening and apoptosis in the pouch and hinge (**Fig 3 A,B**). In contrast to Robo3 overexpression, we found that when we expressed a functional Robo3-RNAi in mosaic clones, we did not observe any interface surveillance responses in the disc (**Fig 3C**, **Fig S3 B,C**). This is consistent with a model where Robo3-RNAi clones do not create pronounced differences in Robo3 levels in an environment that already expresses low levels of Robo3. Combined, we conclude that the expression of a single cell surface molecule, such as Robo3, is completely sufficient to induce all hallmarks of interface surveillance.

**Figure 3.**
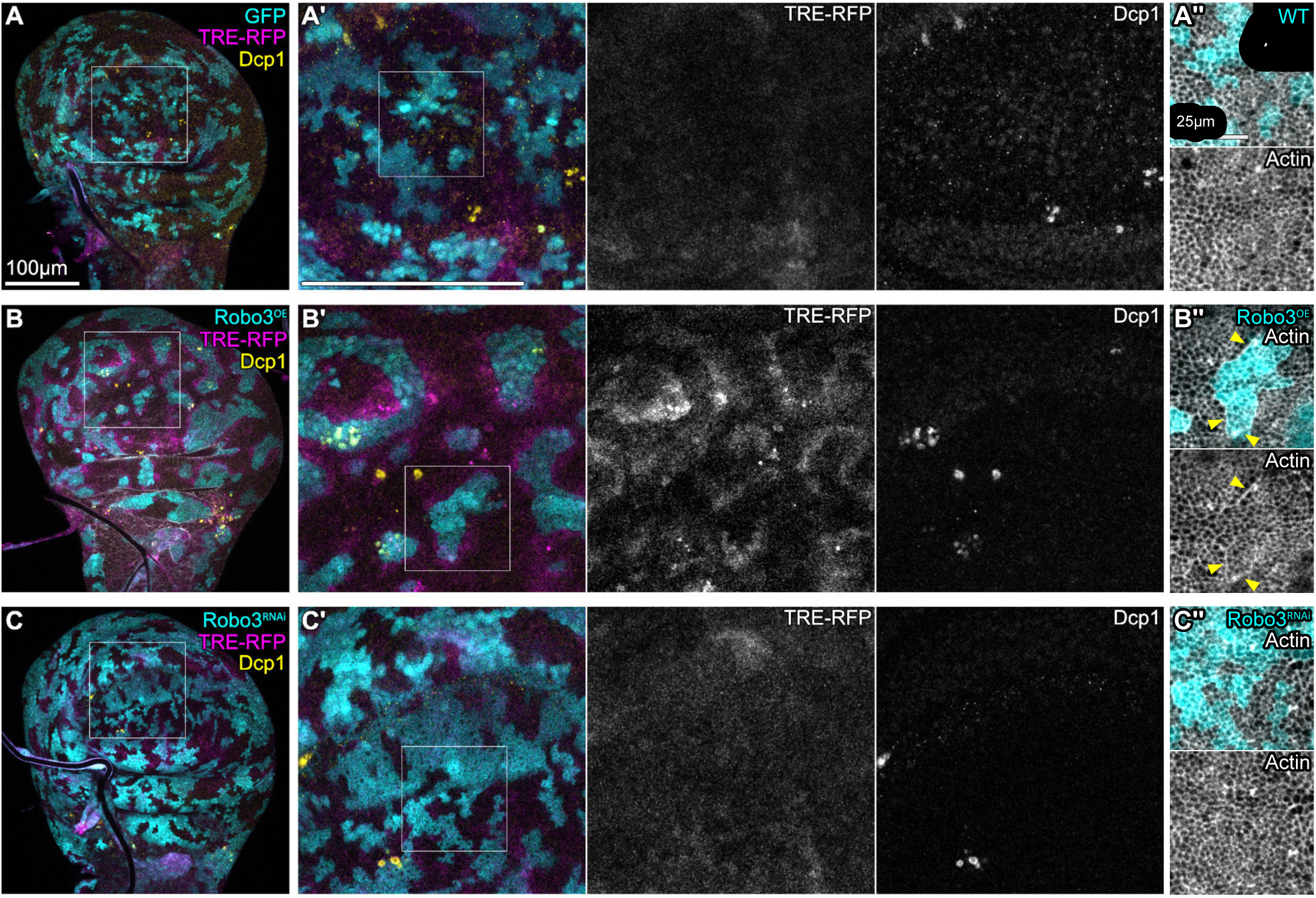
A mismatch of Robo3 receptor expression levels between neighboring cells is detected by interface surveillance in wing disc epithelia. **A** A mosaic wing disc with wild type clones expressing *UAS-GFP* (GFP, cyan). Please note that JNK activity is physiologically elevated at the A/P compartment boundary in wing discs at this developmental stage. **B, C** Mosaic wing discs where clones (cyan) either overexpress *UAS-robo3* (Robo3^OE^) (B), or express *UAS-robo3-RNAi* (Robo3^RNAi^) (C) to reduce Robo3 function. Please note the Actin enrichment at clone interfaces in B’’ (yellow arrows). TRE-RFP expression is reporting JNK pathway activity (grey or magenta in all panels). Antibody staining against the cleaved effector caspase (cDcp1) visualizes apoptosis (grey or yellow). Phalloidin visualizes cortical F-Actin (grey). White frames in (A-C) marks region shown in (A’-C’). White frames in (A’-C’) marks regions shown in (A’’-C’’). Image sets are shown at the same scale. Scale bar in (A’-C’) corresponds to 100µm. Representative wing discs of n ≥ 8 discs of n ≥ 2 experimental replicates.

We next analyzed the effects of mosaic manipulation of Robo2 in the wing disc. Just like Robo3-expressing clones, we find that Robo2-expressing clones have a smooth boundary indicative of actomyosin recruitment to the interface, and induce apoptosis (**Fig 4 D,E**). Moreover, the characteristic pattern of bilateral JNK interface signaling - where both wild type cells and aberrant cells at the clonal interface activate JNK - can be detected around Robo2-expressing clones (**Fig 4 F,H**, **Fig S4 A-C**). JNK interface signaling induced by aberrant cells drives apoptosis to either side of the clone boundary, whereas a less well characterized pathway drives apoptosis in the clone interior ^4^. These two pathways drive apoptosis in both interface cell populations and clone interiors ^4^, a characteristic pattern which could be reproduced by Robo2-expressing clones (**Fig 4I**). Combined these results demonstrate, that misexpression of Robo 2 and Robo 3 can induce all hallmarks of interface surveillance.

**Figure 4.**
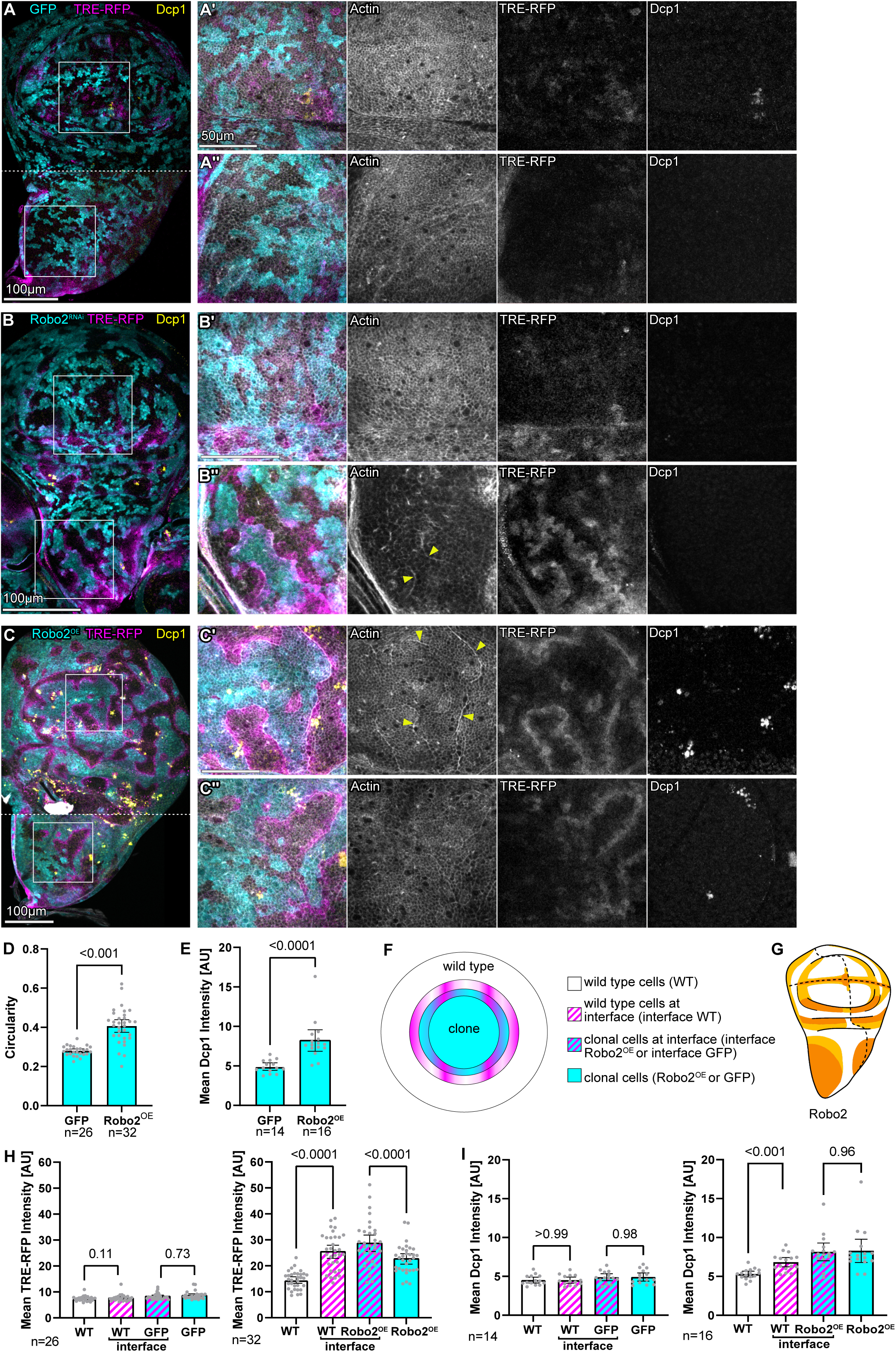
A mismatch of Robo2 receptor expression levels between neighboring cells is detected by interface surveillance in wing disc epithelia. **A** Mosaic wing disc with wild type clones expressing *UAS-GFP* (GFP, cyan). TRE-RFP expression is reporting JNK pathway activity (grey or magenta). Antibody staining against the cleaved effector caspase (cDcp1) visualizes apoptosis (grey or yellow). Phalloidin visualizes cortical F-Actin (grey). Please note that JNK activity is physiologically elevated at the A/P compartment boundary in wing discs at this developmental stage. White frame marks region shown to the right in (A’, pouch and A’’, notum). Dotted line indicates stitching of confocal images with different z-positions. Image sets are shown at the same scale. **B, C** Mosaic wing discs where clones (cyan) either overexpress *UAS-robo2* (Robo2^OE^) (B), or express *UAS-robo2-RNAi* (Robo2^RNAi^) (C) to reduce Robo2 function. TRE-RFP expression is reporting JNK pathway activity (grey or magenta). Antibody staining against the cleaved effector caspase (cDcp1) visualizes apoptosis (grey or yellow). Phalloidin visualizes cortical F-Actin (grey). Please note the Actin enrichment at clone interface (yellow arrows in B’’ and C’). Dotted line indicates stitching of confocal images with different z-positions. White frame marks region shown to the right in (B’, C’, pouch and B’’, C’’, notum). Image sets in (B’-C’’) are shown at the same scale. **D, E** Graphs depicting circularity and apoptosis observed for individual *UAS-GFP* (GFP) and *UAS-Robo2* (Robo2^OE^) expressing clones. Sample number (n) for individual wing discs and p-values of a two tailed, unpaired t-test are displayed in graphs. **F** Illustration of regions of interest (ROIs) relative to the clonal interface defined for quantitative image analysis (see also Fig S4 A-C). There are non-labeled wild type cells (white) or *UAS-GFP* labeled clonal cells (cyan) in the wing disc. Clonal cells which additionally express *UAS-Robo2* are abbreviated as Robo2^OE^. Both, GFP-labeled (cyan) and non-labeled (white) cells are separated into cells which are in contact with the other cell type at the interface (magenta stripes), and the remaining cells away from the interface (see Fig S4 and methods). **G** Illustration of the *robo2-GFP* expression pattern. Relative fluorescence intensity is depicted by color intensity from low (white), medium (light orange) to high (dark orange). **H, I** Graphs depicting mean fluorescence intensity of TRE-RFP reporter and mean cDcp1 intensity in the zones of measurement around clones, as depicted in E. Graphs display results for mosaic discs with *UAS-GFP* labeled clonal cells (left) or containing *UAS-Robo2* (Robo2^OE^) expressing clones (right). Sample number (n) for individual wing discs and p-values of a repeated measured one-way ANOVA with Tukey’s multiple comparisons test are displayed in graphs.

However, in contrast to Robo3, Robo2 is expressed throughout wing disc development, and importantly, in a complex spatial pattern. In third instar discs, Robo2 is most highly expressed in the anterior notum and the hinge-hinge fold region, whereas medium levels of expression can be found along the dorsal-ventral boundary. In the dorsal and ventral pouch, Robo2 expression is low (**Fig 4D**). If Robo2 mediates recognition of cell fate differences, then interface surveillance hallmarks should be activated locally in a Robo2-expression pattern-specific manner by pronounced differences in Robo2 expression levels between neighboring cells. Therefore, we predicted that Robo2-expressing clones should elicit interface surveillance responses in the pouch but not the anterior notum, whereas Robo2-RNAi expressing clones should elicit pattern-specific responses in the notum and not the pouch. Indeed, Robo2-expressing clones induced Actin enrichment and JNK interface signaling in the pouch but not the notum, whereas Robo2-RNAi expressing clones induced this set of responses in the notum but not the pouch (**Fig 4 A-C**). These spatially restricted responses for Robo2-expressing and Robo2-RNAi expressing clones which depend on endogenous Robo2 expression patterns can be visualized in a Robo2-GFP expressing disc (**Fig S4 D,E**). These results demonstrate that cells which create pronounced differences in Robo2 expression levels are robustly detected by interface surveillance within the endogenous Robo2 expression pattern. Combined, these experiments demonstrate the common pattern-specific function of Robo2 and Robo3 receptors in inducing interface surveillance, as previously shown for fate-specifying transcription factors expressed in different spatial patterns. We thus suggest that these cell surface molecules mediate cell fate recognition in epithelial tissues and thereby contribute to the maintenance of epithelial health by initiating detection and removal of aberrant cells.

### Expression of Robo receptors and other cell surface molecules in wing disc epithelia is regulated by fate-patterning pathways

If cell surface molecules establish cell recognition in interface surveillance, then interface surveillance-inducing aberrations of patterning pathways should alter the expression pattern of these proteins to create pronounced expression level differences. To test this, we created clones misexpressing regulators of different fate-patterning pathways: Thickveins (*tkv^CA^*), a constitutively active Dpp/TGFβ-receptor. Cubitus interruptus (*ci*), the transcription factor activated by Hedgehog (Hh)-signaling. Armadillo (*arm^S10^*), a constitutively active co-activator of the TCF/LEF effectors of Wg/Wnt signaling. Eyeless (*ey*), a Pax6 homologue in the eye-specification network. Forkhead (*fkh*), a transcription factor required for salivary gland specification. We then tested how targeted expression in clones may alter expression of interface surveillance-inducing cell surface molecules (Robo2, Eph, or Ten-m; **Fig 5 A-C**).

**Figure 5.**
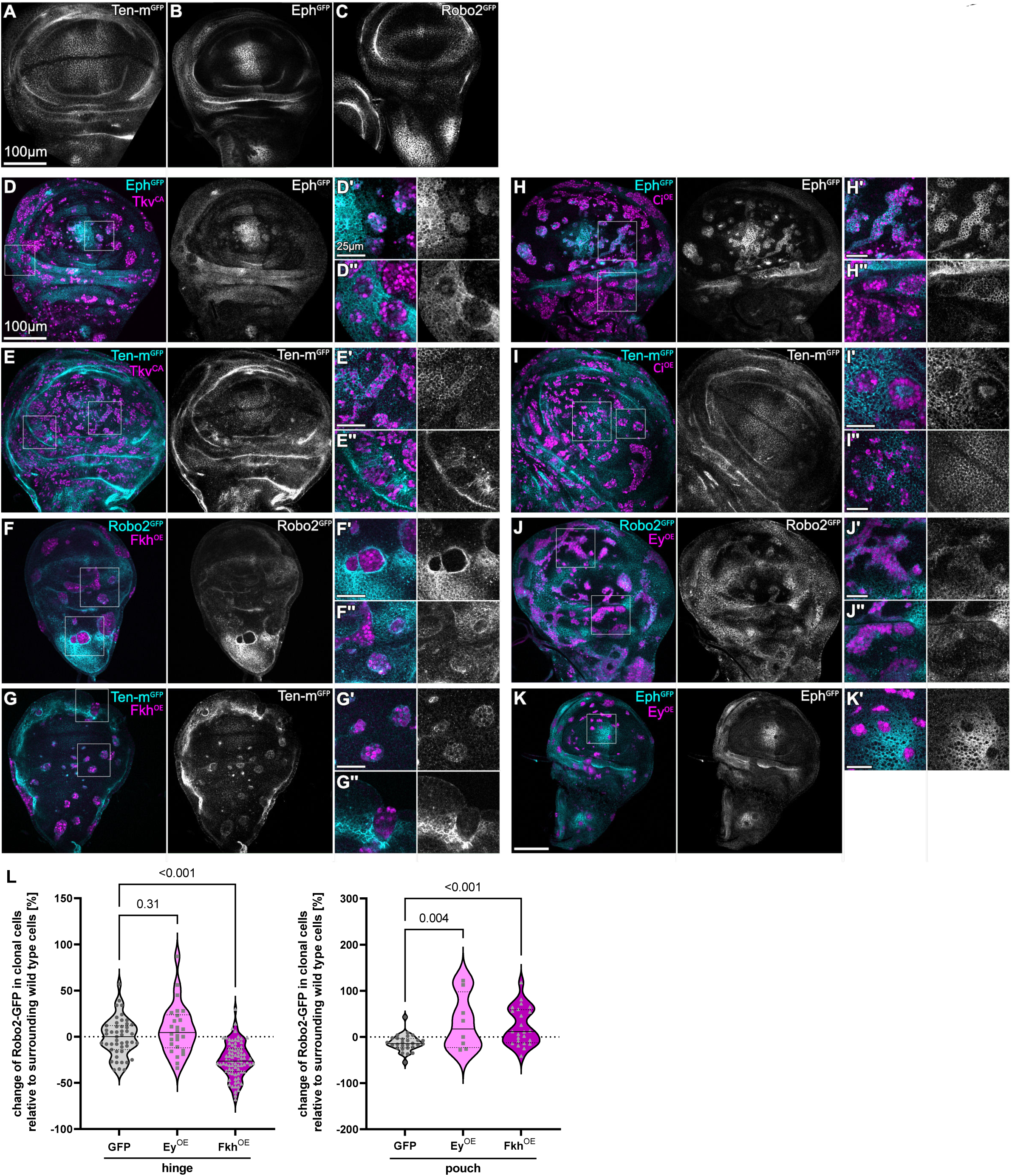
Expression of Robo receptors and other cell surface molecules in wing disc epithelia is regulated by fate-patterning pathways. **A-C** Wing discs presenting the specific expression patterns of *ten-m-GFP* (Ten-m^GFP^) (A), *eph-GFP* (Eph^GFP^) (B) and *robo2-GFP* (Robo2^GFP^) (C) and serve as a reference for observed changes in (D-K). **D-K** Wing discs expressing *eph-GFP* (Eph^GFP^), *robo2-GFP* (Robo2^GFP^) or *ten-m-GFP* (Ten-m^GFP^) fusion proteins (grey or cyan). Mosaic clones (magenta) deregulate fate specifying pathway by expression of a constitutively active Tkv (Tkv^CA^) receptor (D,E), Fkh (Fkh^OE^) (F,G), Ci (Ci^OE^) (H,I), or Ey (Ey^OE^) (J,K). White frames mark regions shown in (D’-K’ and D’’-J’’). **L** Graphs depicting change of *robo2-GFP* in clonal cells expressing *UAS-GFP, UAS-ey or UAS-fkh* relative to surrounding wild type cells in the hinge or pouch region of the wing disc. Clones of ≥5 wing discs per genotype are plotted and p-values of a one-way Anova with Dunnett’s multiple comparison test are displayed in graphs. For *robo2-GFP* genotypes, n=8 wing discs in n=3 experimental replicates were analyzed in J and n = 22 wing discs in n = 2 experimental replicates in F. Image sets are shown at the same scale.

Strikingly, we found that expression patterns of these molecules were significantly altered and that, importantly, manipulation of a single-fate-specifying pathway regulated multiple molecules (**Fig 5**, **Fig S5**). For example, *tkv^CA^*-expressing clones altered expression of Eph, Ten-m and Ten-a (**Fig 5 D,E**, **Fig S5F**). Yet, these same molecules, as well as Robo 2, were also regulated by manipulating other fate-specifying pathways, such as misexpression of *ci*, *fkh* or *ey* (**Fig 5 F-L**, **Fig S5 A-D,G,H**). Adding to the complexity, we observed that regulation depended on the position of clones in the disc. For example, *tkv^CA^*-expression caused Eph upregulation in the pouch but downregulation in the hinge of the disc (**Fig 5 D’,D’’**). These findings emphasize that one fate-specifying pathway does not exclusively upregulate (or downregulate) expression of a specific cell surface molecule. Rather, multiple developmental patterning pathways are integrated to drive expression of a specific set of genes at any spatial position of the wing imaginal disc. Consequently, highly cell-type specific decisions determine whether to express a target gene encoding a cell surface molecule or not. This conclusion is consistent with patterning models of the disc ^40, 46^.

### Interface surveillance must be mediated by multiple cell surface receptors

We found that many cell surface molecules induce interface surveillance responses, and alteration of a single fate program changes expression of multiple cell surface molecules. We thus predicted that genetic rescue experiments targeting a single molecule may not be sufficient to interfere with interface surveillance responses to differently fated cells. To test this, we expressed UAS-RNAi for molecules that are upregulated in the peripheral pouch and hinge domain by *tkv^CA^* (where Tkv is not normally active) under the control of *en*-GAL4. Thereby, all cells in the posterior compartment lacked either Eph or Ten-m at the time when we introduced *tkv^CA^*-expressing clones using the LexA/LexO system ^47, 48^. However, removing Eph or Ten-m from the surface code failed to affect interface surveillance responses of aberrant *tkv^CA^*-expressing clones in the posterior pouch and hinge (**Fig 6**, **Fig S6 A-D**). Similarly, other aberrant cell types induced by *ey, fkh,* or *ci*-expression still induced enrichment of Actin at clonal interfaces, clone smoothening and even apoptosis, despite intra-clonal rescue experiments to restore expression levels of the cell surface molecules Eph, Robo2 or Ten-m (**Fig S6 E-H**). These experiments demonstrate that the induction of interface surveillance by aberrantly positioned cells is likely caused by deregulation of more than just one cell surface molecule. We conclude that developmental patterning pathways regulate the expression of multiple cell surface molecules and that, while deregulation of one cell surface molecule like Robo2 is sufficient, alteration of these multiple molecules in clones with inappropriate cell fate causes highly redundant recognition and activation of interface surveillance.

**Figure 6.**
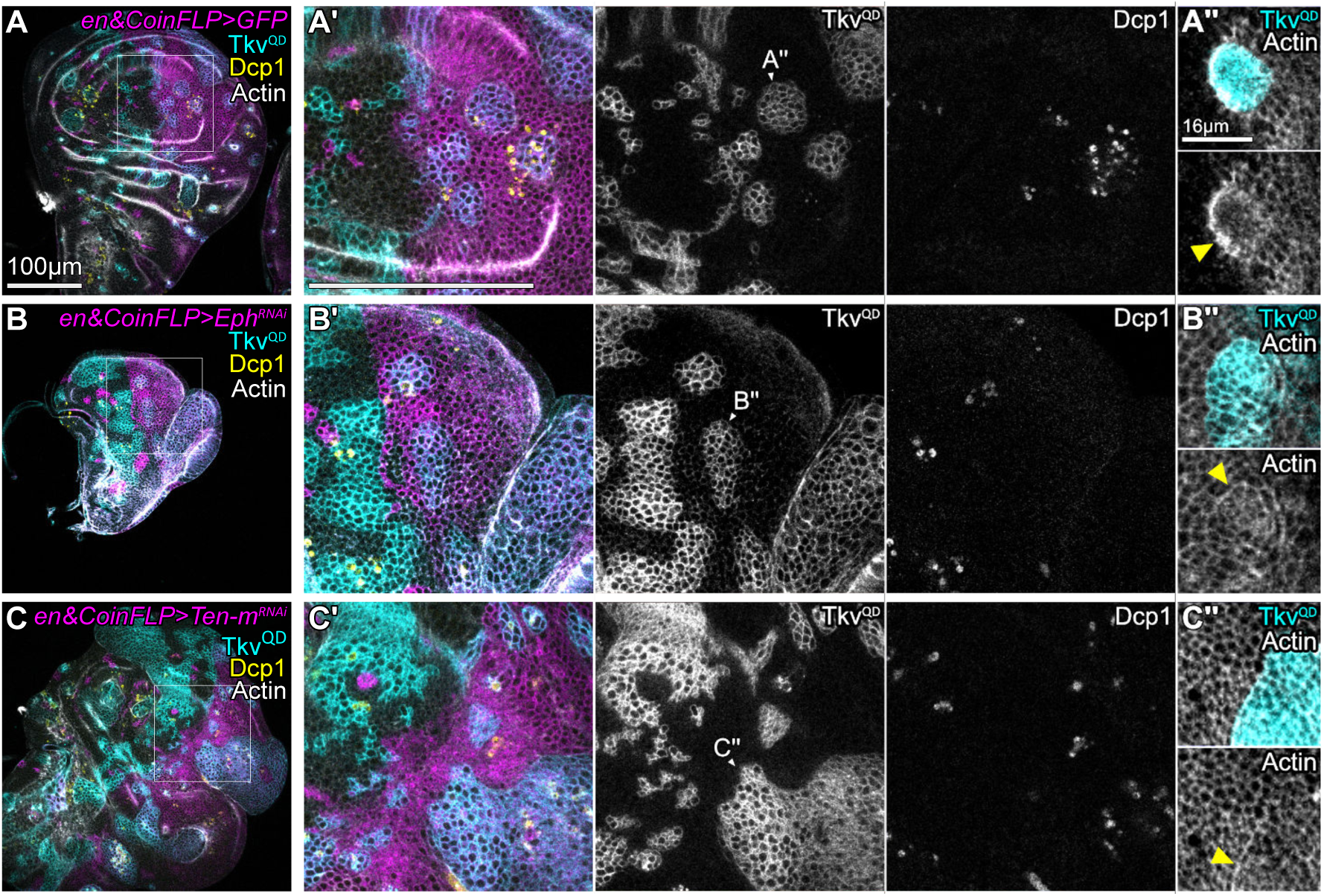
Interface surveillance is redundantly mediated by multiple cell surface receptors. **A-C** Mosaic wing discs with clones deregulating a fate-specifying pathway by expressing a constitutively active Tkv receptor (Tkv^QD^) via the LexA/LexO-CoinFLP system (grey or cyan). *en*-GAL4 constitutively drives expression of UAS-constructs in the posterior compartment (magenta). The CoinFLP induces either LexA or GAL4 clones upon Flp-FRT recombination. Therefore GAL4/UAS clones expressing *UAS-GFP* (GFP) can also occasionally be observed in the anterior compartment (magenta, see also Fig S6 B,D). All GAL4-expressing cells (magenta) drive knock-down of cell surface molecules by expression of *UAS-eph-RNAi* (Eph^RNAi^) (B) *or UAS-ten-m-RNAi* (Ten-m^RNAi^) (C) (see Fig S6 A,C). Antibody staining against the cleaved effector caspase (cDcp1) visualizes apoptosis (grey or yellow). Phalloidin visualizes cortical F-Actin (grey). White frames mark regions shown in (A’-C’). Images are shown at the same scale in (A-C) and (A’-C’). **A’’-C’’** Magnified view of clones in the posterior compartment marked in (A’-C’). An apical section is shown. Please note the Actin enrichment at clonal interfaces (yellow arrows). Images are shown at the same scale. Representative wing discs of n ≥ 5 wing discs from n = 2 experimental replicates are shown.

### Oncogenic Ras but not classical cell-cell competition alters expression of the cell surface code

To better understand how regulation of cell surface receptors may be relevant for pathological processes, we turned to the analysis of oncogenic mutations incurred by *Ras^V12^*. Ras-signaling is a component of the EGF/ERK signaling axis and affects gene expression, for example via activation of ETS transcription factors. The pathway is tightly controlled to regulate proliferation, but also vein cell fate patterning in late larval and pupal wing tissue ^49^. We previously reported that *Ras^V12^*-expressing cells elicit interface surveillance in imaginal discs, as evident by enrichment of Actin at clone boundaries, clone smoothening and activation of JNK interface signaling (**Fig 7A**) ^3, 50^. Yet, *Ras^V12^*-expressing cells evade elimination by apoptosis because of the potent anti-apoptotic properties conferred by *Ras^V12^*-expression, thereby allowing these cells to remain in the tissue and form slow-growing tumors ^4^. Thus, while *Ras^V12^*-expressing cells may not be eliminated by interface surveillance mechanisms, the surrounding wild type tissue still recognizes them as aberrant cells as part of the interface surveillance detection system.

**Figure 7.**
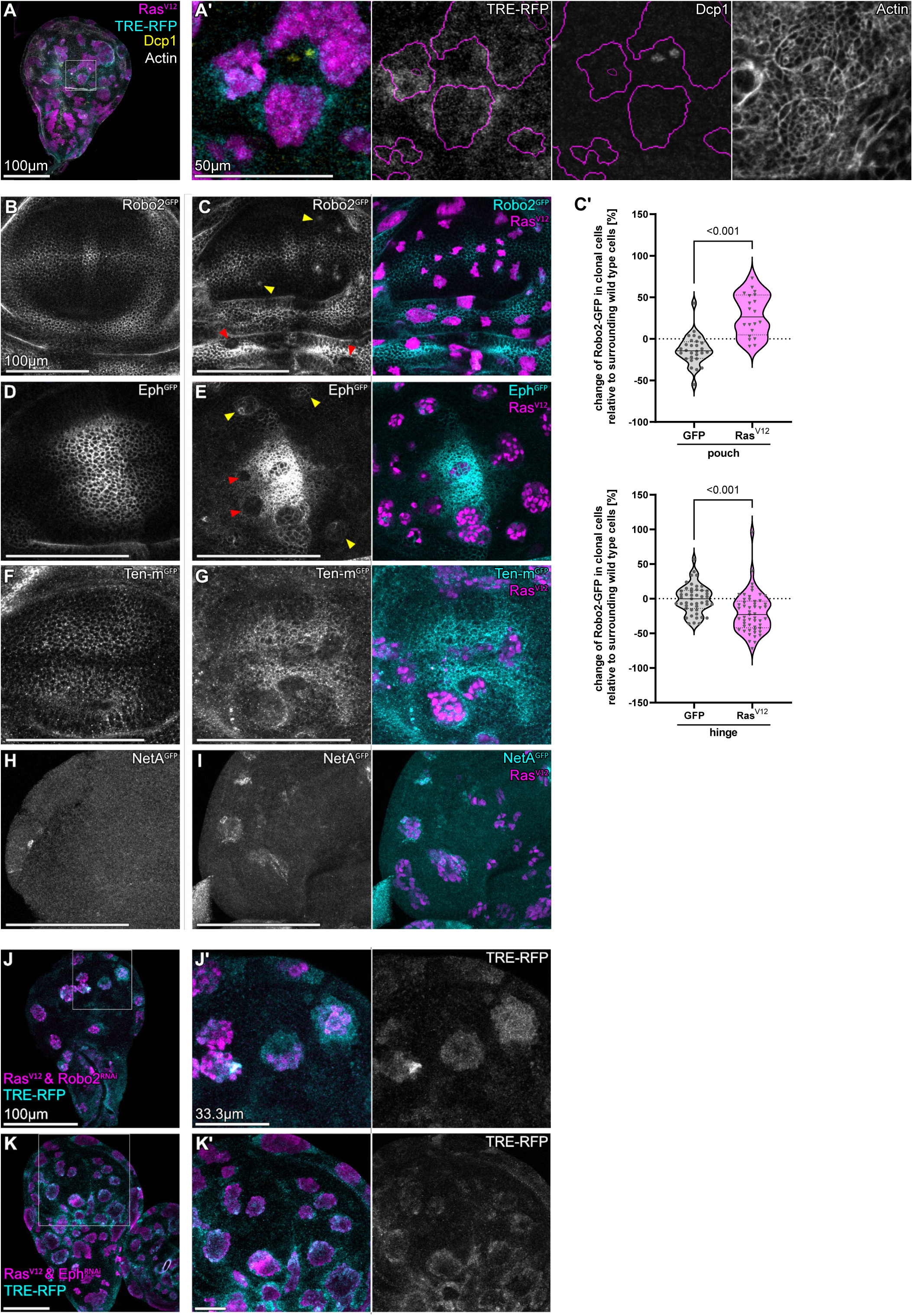
Oncogenic Ras alters expression of the cell surface code. **A** Mosaic wing discs with clones deregulating EGF/ERK-signaling by expression of an oncogenic Ras, using the *UAS-RasV12* (Ras^V12^) (magenta) construct. TRE-RFP expression is reporting JNK pathway activity (grey or cyan). Antibody staining against the cleaved effector caspase (cDcp1) visualizes apoptosis (grey or yellow). Phalloidin (grey) visualizes cortical F-Actin. White frame marks region shown in (A’). Outline of clonal marker in magenta. **B,D,F,H** Wing discs presenting the specific expression patterns of *robo2-GFP* (Robo2^GFP^) (B), *eph-GFP* (Eph^GFP^) (D), *ten-m-GFP* (Ten-m^GFP^) (F) or *netA-GFP* (Net^GFP^) (H), serving as a reference for the experimentally induced changes in (C,E,G,I). Image sets are shown at the same scale. **C,E,G,I** Wing discs expressing *robo2-GFP* (Robo2^GFP^) (C), *eph-GFP* (Eph^GFP^) (E), *ten-m-GFP* (Ten-m^GFP^) (G) or *netA-GFP* (Net^GFP^) (I) (grey or cyan) and carrying mosaic clones (magenta) expressing *UAS-RasV12* (Ras^V12^). Yellow arrows point to clones with elevated expression of *robo2-GFP* (Robo2^GFP^) (C) or *eph-GFP* (Eph^GFP^) (E), red arrows point to clones with reduced expression of *robo2-GFP* (Robo2^GFP^) (C) or *eph-GFP* (Eph^GFP^) (E). Image sets are shown at the same scale in (B-I). **C’** Graphs depicting change of *robo2-GFP* in clonal cells expressing *UAS-GFP or UAS-Ras^V12^* relative to surrounding wild type cells in the hinge or pouch region of the wing disc. Clones of ≥5 wing discs per genotype are plotted and p-values of a one-way Anova with Dunnett’s multiple comparison test are displayed in graphs. Please note that UAS-GFP control clones are the same as in Fig 5. **J,K** Mosaic wing discs with clones expressing oncogenic *UAS-RasV12* (Ras^V12^) (magenta). The same clones either co-express *UAS-robo2-RNAi* (Robo2^RNAi^) (J) or *UAS-eph-RNAi* (Eph^RNAi^) (K). TRE-RFP expression is reporting JNK pathway activity (grey or cyan). White frames marks regions shown in (J’,K’). Please note that clones still acquire a round smooth shape. Image sets are shown at the same scale in (J,K) and (J’,K’).

To understand, if *Ras^V12^*-expressing cells are initially faithfully detected by altered expression of cell surface markers, we monitored expression of multiple cell surface molecules in *Ras^V12^*-expressing mosaic disc. Indeed, *Ras^V12^*-expressing clones alter expression of Robo2, Eph, Ten-m, Net A and others (**Fig 7 B-I**, **Fig S7**). Importantly, *Ras^V12^* alters the expression of these molecules depending on the spatial position in the disc. For example, *Ras^V12^* mildly elevated expression of Robo2 in the pouch but downregulated Robo2 in the hinge domain (**Fig 7C**). This confirms our results above that alterations in a single pathway cooperate with cell-type specific patterning information to control expression of genes encoding cell surface molecules. To also confirm that restoring expression of a single cell surface molecule in aberrant cells is not sufficient to prevent detection by interface surveillance, we designed a rescue experiment where we co-expressed an RNAi-construct targeting either Robo2 or Eph in *Ras^V12^*-expressing clones. We specifically aimed to removed Eph or Robo2 expression in clones that ectopically induce expression of Eph or Robo2 in the peripheral pouch. However, these clones still activated interface surveillance responses (**Fig 7 J,K**), again strongly supporting our conclusion that in *in vivo* expression alterations in more than one cell surface receptor can mediate detection of *Ras^V12^*-expressing cells.

Finally, we wanted to understand if other established tissue-intrinsic error-correction mechanisms, such as classical cell competition, may utilize the same cell surface code to distinguish loser and winner fates. In cell competition, the comparison of proteostatic or metabolic fitness between neighboring cells drives cell elimination, such that less fit ’loser’ cells are eliminated by fit ’winner’ cells ^51–58^. Previous studies indicate that loser cells start to express a specific isoform of the cell surface molecule Flower, which marks them for elimination by winner cells in a cell-contact dependent manner ^59, 60^. To test if also a Robo2, Robo3, Eph or Ten-m-dependent cell surface code is associated with winner or loser cell state, we analyzed expression of these cell surface molecules in imaginal discs mosaic for a Myc-expressing winner genotype. In agreement with the model that cell competition and interface surveillance are mechanistically distinct error correction mechanisms, we found that Myc-expressing clones did not display pronounced alterations of cell surface receptor expression patterns that were predictive of either winner or loser fate (**Fig 8**). Moreover, Myc-expressing clones do not induce interface tension in mosaic tissues ^61^. Similarly, other genotypes linked to cell competition, tissue growth or cell survival, such as moderate activation or inhibition of Hippo/Warts or STAT92E signaling, did not alter cell surface molecule expression patterns (**Fig S8**). These observations suggest that a change in expression patterns of cell surface molecules is specific to cell fate specifying pathways and detection by the interface surveillance program. This conclusion is strongly supported by our previous studies, in which cell competition and interface surveillance form distinct branches of tissue-intrinsic error-correction and tumor suppression mechanisms that maintain the health of epithelial tissues ^4^.

**Figure 8.**
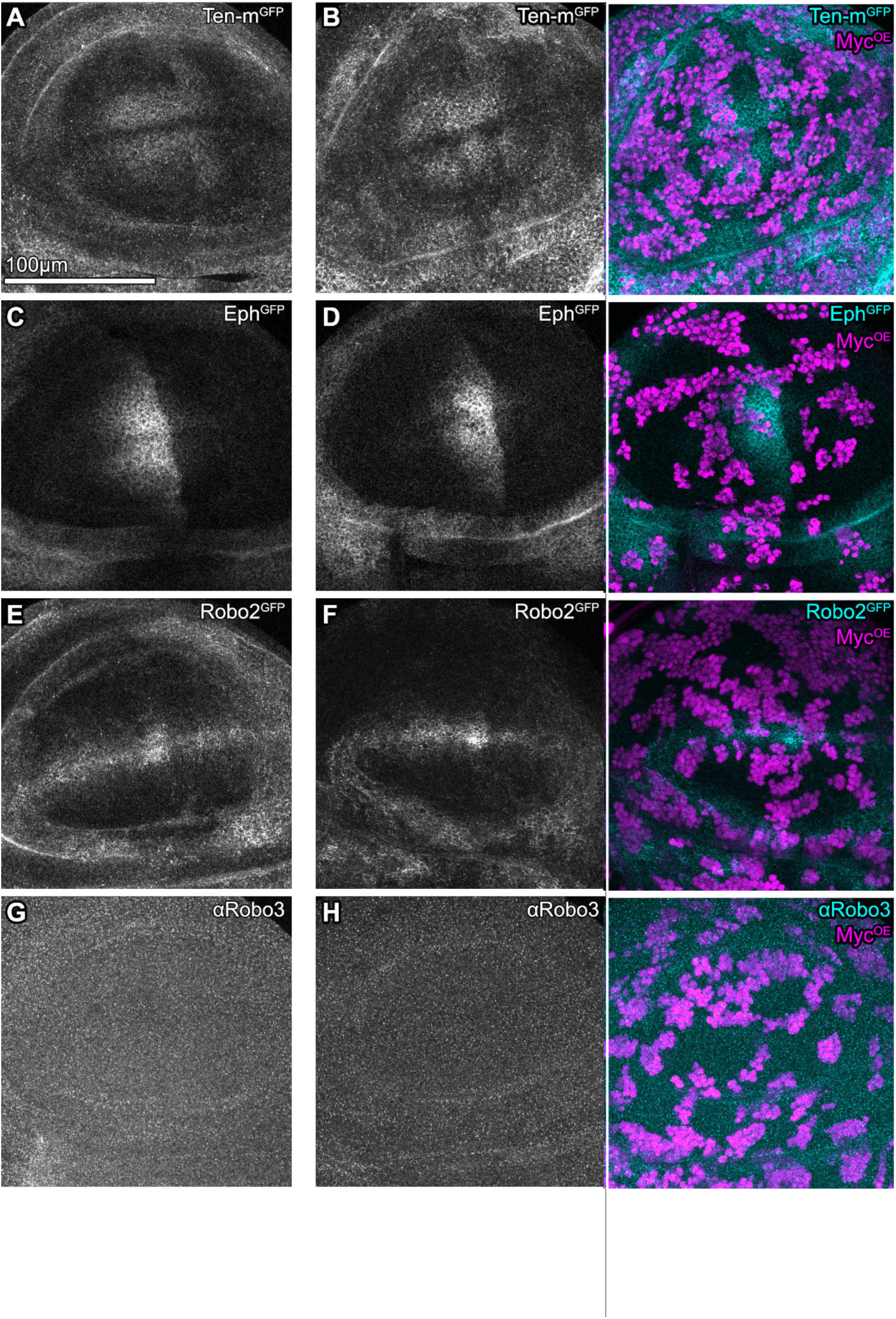
Classical models of cell-cell competition do not regulate the cell surface code. **A,C,E,G** Wing discs presenting the specific expression patterns of *ten-m-GFP* (Ten-m^GFP^) (A), *eph-GFP* (Eph^GFP^) (C)*, robo2-GFP* (Robo2^GFP^) (E) or stained for Robo3 (G) serving as a reference for the experimentally induced changes in (B,D,F,H). **B,D,F,H** Wing discs expressing *ten-m-GFP* (Ten-m^GFP^) (B), *eph-GFP* (Eph^GFP^) (D)*, robo2-GFP* (Robo2^GFP^) (F) or stained for Robo3 (H) (grey or cyan) and carrying mosaic clones which overexpress Myc (Myc^OE^) (magenta). Increasing Myc-levels creates super competitor cells, which will outcompete surrounding wild type cells. Images are shown at the same scale.

## Discussion

### Cell fate, cell surface code and interface surveillance

In this study we demonstrate that distinct cell surface molecules are regulated by cell-fate-specifying transcription factors and developmental patterning pathways. Because each pathway can target multiple cell surface molecules, cell fates can establish a complex cell surface code that is unique to each fate. A mismatch in the cell surface code created by changes in cell-fate-specifying transcription factors and developmental patterning pathways then induces local responses associated with interface surveillance in a highly redundant manner (**Fig S9.1**). Importantly though, creating pronounced differences in expression levels of a single surface molecule is sufficient for cells at the expression boundaries to respond with interface Actin recruitment, clone smoothening and bilateral JNK interface signaling.

Of the protein families that we identified in our genetic screen, specifically Eph/Ephrin signaling has been previously described to promote interface smoothening via actomyosin regulation in mosaic tissues and at compartment boundaries, both in Drosophila and mammalian systems ^36, 43^. Indeed, deregulation of Eph/Ephrin induces actomyosin enrichment at clonal interfaces, clone smoothening and bilateral JNK signaling in our assays (**Fig S9.2**). Similarly, Teneurins, LRR proteins and atypical Cadherins have been implicated in modulating actomyosin contractility or compartment boundary smoothening in different tissues ^36, 37, 44, 61–64^. Yet, we also describe a novel function of these molecules in inducing bilateral JNK activation. This suggests that these very different cell surface molecules may act not only through a shared set of cytoskeleton effectors but also through similar signaling effectors. The cell surface molecules identified in our screen are composed of very different structural motifs, including leucine-rich, immunoglobulin, YD or cadherin repeats ^29, 30, 35, 65^. How these extracellular motifs or intracellular domains contribute to these shared effector modalities of actomyosin contractility and JNK signaling remains to be determined.

### Robo receptors in epithelia

Robo receptors are increasingly recognized to play a role in epithelia ^29^. For example, recent molecular data obtained from the mouse pancreas showed that epithelial islet cells express Robo receptors. Inactivating Slit-Robo signaling disrupted embryonic islet formation and adult islet architecture ^66, 67^, emphasizing an underappreciated role for axon guidance molecules in extra-neuronal contexts. In Drosophila, Robo2 was suggested to be induced by high cell-autonomous JNK signaling and required for the elimination of *scrib* mutant cells ^45^. How these observations are related remains to be investigated.

From our genetic screen for interface surveillance molecules, we specifically isolated Robo2 and Robo3. Robo2 and 3 are closely related molecules with previously described function in neuronal cell-cell recognition, and are different from Robo1. While all Robo receptors share similar extracellular domains, Robo2 and 3 retain only two of the four cytoplasmic proline-rich CC-domains found in Robo1 and lack the Ena-recruiting motif. Curiously, all Robo receptors retain a WAVE Regulatory Complex binding site through which Actin regulation may be facilitated ^68, 69^. As described above, altering expression levels of Robo2 induced interface surveillance in a Robo2-pattern-specific manner everywhere in the disc. Yet, Robo3 displays one exception to the expected pattern-specific response: Robo3-expressing clones did not induce the expected interface surveillance responses in a small region of the proximal, anterior notum. Instead, the response patterns resembled that of Robo2 (**Fig S3 D,E**). We suggest that Robo2 and Robo3 may interact in similar receptor complexes, yet their relationship needs to be explored in future studies.

### Regulation of neuronal cell surface molecules in cancer

Multiple recent studies revealed significant changes to the expression patterns of neuronal cell surface molecules in tumors for example ^70, 71^. We demonstrate that Ras^V12^-expressing cells disrupt the endogenous pattern of surface molecules, causing pronounced differences in the surface code and activation of interface surveillance responses. In mammalian epithelia, cells with oncogenic Ras signaling are detected and eliminated by a process termed EDAC akin to interface surveillance ^4, 72, 73^. Eph/Ephrin signaling was shown to regulate the interaction of oncogenic Ras^V12^-expressing cells with neighboring wild type cells in cultured MDCK monolayers and pancreas *in vivo* ^74, 75^. Here, Ras^V12^-expression induces Eph2A which underlies Actin enrichment and cell shape changes upon interaction of wild type and Ras^V12^-expressing cells. This indicates that interface surveillance mediated by cell surface molecules may have important roles in tumor suppression by regulating the interaction between wild type and aberrant cells. Yet, the same machinery may also facilitate tumor progression by activating signaling and actomyosin dynamics at the interface to aberrant cells, which may be resistant to apoptosis and therefore to elimination, as observed for Ras^V12^-expressing cells ^4^.

### Interface surveillance in development

Interface surveillance shares striking similarities with compartment boundary formation in developing tissues, where two cell populations of distinct fate mechanically segregate via the formation of a contractile actomyosin interface between them. Indeed, many of the molecules identified in our genetic screen facilitate actomyosin regulation during compartment boundary formation in vertebrate and invertebrate species ^24–26, 36, 37^. The similarities between developmental morphogenesis and tissue-intrinsic error-correction, relying on the same principle of receptor-mediated recognition and actomyosin driven segregation of cell populations, point towards a common evolutionary origin of the surface code subsequently adapted for developmental and homeostatic processes. In fact, cellular function of neuronal axon guidance in neuronal tissues may have originally evolved in evolutionarily more ancient epithelia. Ultimately, the cell surface code formed by so-called neuronal axon guidance molecules and related factors may represent an ancient system to distinguish self from non-self in physiological and pathological contexts.

## Experimental procedures

### *Drosophila* stocks and genetics

*Drosophila melanogaster* fly strains (**Supplementary Table S1** and **Supplementary Table S2**) were maintained on standard fly food (10L water, 74,5g agar, 243g dry yeast, 580g corn flour, 552ml molasses, 20.7g Nipagin, 35ml propionic acid) at 18°C - 22°C. Larvae from experimental crosses were allowed to feed on Bloomington formulation (175.7g Nutry-Fly,1100ml water 20g dry yeast, 1.45g Nipagin in 15ml Ethanol, 4.8ml Propionic acid) and raised at 25°C. Mosaic gene expression was induced by Flip-out (act or tub>GAL4/UAS, CoinFLP-LexA/LexO) or mitotic FLP/FRT ^42, 47^. Parental adult flies were allowed to lay eggs for 72 h at 25°C before flippase expression was induced by heat shock (HS) at 37°C for 5-15 min. Detailed genotypes and experimental conditions are listed in **Supplementary Table S3**. For the JNK interface signaling screen (**Fig S2.1F**), larvae were dissected 48 h after a 10 min HS (AHS). Due to genetic limitations, the CoinFLP experiments (**Fig 6**) contained two GAL4-drivers, the Actin5C-FRT/STOP-GAL4 and the *en*-GAL4, resulting in mosaic GAL4 expression in the total disc and GAL4 expression in the posterior compartment. For experiments examining expression patterns (**Fig 1 and S1**), larvae were dissected 80 h or 102 h after egg lay (AEL). CoinFLP control experiments (**Fig S6 A,C**) were dissected at 102 h AEL.

### Immunohistochemistry

Larvae were dissected and the inverted cuticula with attached wing discs were fixed in 4% paraformaldehyde in PBS for 15 min at 22°C. Samples were washed in 0.1% Triton X-100 in PBS (PBT) and incubated with primary and secondary antibodies in 400µl PBT on a nutator overnight at 4°C and 1-3 h at 22°C, respectively. Following a final wash in PBS, wing discs were mounted with Molecular Probes Antifade Reagents (#S2828). Following antibodies were used: Mouse anti-Robo1 (1:20, DSHB-13C9), mouse anti-Robo3 (1:20, DSHB-14C9&15H2), mouse anti-Ds (1:1000, Suzanne Eaton), mouse anti-Nrt (1:20, DSHB-BP106), mouse anti-Sema2 (1:20, DSHB-19C2), mouse anti-Ptc (1:20, DSHB-apa1), mouse anti-Wg (1:100, DSHB-4D4), rabbit anti-Ephrin (1:1000, Andrea Brand), rabbit anti-Dcp1 (1:400, Cell Signaling-9578S), rat anti-E-cadherin (1:50, DSHB-DCAD2), mouse anti-1-gal (1:1000, Promega Z378B), mouse anti-1-gal (1:1000, MP Biochemicals-55976), rabbit anti-GFP (1:400, Thermo Fisher-G10362), rat anti-RFP (1:20, Heinrich Leonhardt, 5F8) and rat anti-RFP (1:2000, ChromoTek-5F8). Fluorescent dyes DAPI (0.25 ng/µl, Sigma), Phalloidin-Alexa Fluor 405 (1:2000, Abcam-ab176752), Phalloidin-Alexa Fluor 488 (1:500, Invitrogen A12379), Phalloidin-Alexa Fluor 555 (1:500, sigma-P1951) and Phalloidin-Alexa Fluor 647 (1:500, Invitrogen A22287) were added together with secondary antibodies.

### Image acquisition and image display

Samples were imaged using Leica TCS SP8 confocal microscopes. Figure panels were assembled in Affinity Designer. In most figure panels, individual channels represent the same z-position from a confocal image stack. However, some panels may assemble channels from different z-positions in the wing disc (as listed in **Supplementary Table S4**). Such a portrayal was chosen to better visualize the spatially distinct phenotypes at different positions within the cell, specifically of (1) junctional actin (apical), (2) cytoplasmic/nuclear RFP (lateral) and (3) cDcp1 (basal) in one image panel. This was meant to reduce the (peripheral) data load in the manuscript that would be otherwise required to visualize all channels at all positions.

If mentioned in the figure legends, the local-z-projector Fiji plugin (https://gitlab.pasteur.fr/iah-public/LocalZprojector, v1.5.4) ^76^ was used to project the curved apical surface of the wing disc epithelium into the same plane. The local-z-projector generates a surface height-map based on a reference channel (i.e., Ecad or Phalloidin staining) and other channels of interest were displayed in relation to this surface height-map.

### Image analysis, quantification and statistics

To provide a measure for reproducibility and robustness, number of samples (wing discs) and experimental replicates are mentioned in the figure legends. An experimental replicate refers to an independent event for dissection and sample processing. From each larva, only one wing disc per larva was dissected and considered to be an independent sample for statistical analysis. Please note that several discs were visually analyzed for each experimental replicate. For all quantifications, control and experimental samples were processed together and imaged in parallel, using the same confocal settings. For comparison of TRE-RFP and cDcp1 fluorescence intensities in different bands around the clone, a repeated measured one-way ANOVA with Tukey’s multiple comparisons test was used. One-way anova with Dunnett’s multiple comparison test was used for comparison of changes in Robo2-GFP intensities in aberrant clones. For other comparisons, a two tailed, unpaired t-test was performed. Images were processed in FIJI ^77^. GraphPad Prism (version 9.5.0) was used for statistical analysis and generation of graphs.

### Region of interest {ROI) segmentation and quantification workflow

(ROI1) of total disc area: RFP or Phalloidin signal was thresholded and the ROI created by Convert to Mask, Fill Holes, and selection by the wand tool. (ROI2) of clone area: clonal GFP signal was thresholded and ROIs were created by analyze particles=10µm-infinity. For cDcp1 quantification, the lower cutoff was 2µm. (ROI3) WT cells at interface: the Make Band, 4µm command created a band around the ROI2. (ROI4) clonal cells not at the interface: the Enlarge, -4µm command applied on ROI2 created a ROI of the inner clone area. (ROI5) Clonal cells at the interface: XOR of ROI2 and ROI4. (ROI6) WT area around the ROI3: First, Make Band, 16µm created a band around ROI2. XOR of this band and ROI3 resulted in ROI6.

Subsequent general procedures: by using the Combine command, several ROIs of a category, such as individual clones in ROI2, were combined to one ROI and used for further analysis. Signal outside the wing disc was excluded by AND of ROI1 and other ROIs. ROIs of WT cells, such as ROI3 and ROI6, might cover clonal area due to insufficient spacing of clones. This was prevented, in example for ROI3, by first selecting OR of ROI3 and ROI2, followed by XOR of that selection and ROI2. After the disc segmentation into ROIs, the Measure command was applied.

#### Workflow for quantification of TRE-RFP intensity in UAS-eph-RNAi and UAS-GFP clones

7min HS at 37°C, 72 h AEL, reared at 25°C, dissected 72 h AHS. TRE-RFP signal was elevated by anti-RFP antibody staining. One lateral section was imaged and quantified. Area and mean intensity of TRE-RFP was measured in each ROI. For circularity, the shape of individual clones in ROI2 was measured.

#### Workflow for quantification of TRE-RFP intensity in UAS-robo2-HA and UAS-GFP clones

8min HS at 37°C, 72 h AEL, reared at 25°C, dissected 48 h AHS. TRE-RFP signal with and without rat anti-RFP antibody was compared (**Fig S4**). One lateral section of native TRE-RFP was imaged and quantified. Area and mean intensity of TRE-RFP was measured in each ROI. For circularity, the shape of individual clones in ROI2 was measured.

#### Workflow for quantification of Dcp1 intensity in UAS-robo2-HA and UAS-GFP clones

9min HS at 37°C, 72 h AEL, reared at 25°C, dissected 30 h AHS. In total, 4 confocal z-sections were quantified. As apoptotic cells are localized basally, the stack was started at the most basal position, moving towards the apical surface in 2µm steps, resulting in the quantification of basal and lateral sections. The above-described ROI segmentation and measurement of cDcp1 intensity was applied individually for each z-section in all ROIs. Eventually, the intensities of all four z-sections were averaged before doing the statistics.

#### Workflow for quantification of Robo2-GFP regulation in UAS-GFP, UAS-ey, UAS-RasV12, UAS-fkh clones

10min HS at 37°C, 72 h AEL, reared at 25°C, dissected 44 h AHS. Confocal z-stack had a step size of 4µm. Of those, 4 contiguous lateral z-sections beneath the peripodium were chosen for quantification. First, the wing disc was manually separated into approximate pouch, hinge and notum areas by using the polygon selections in Fiji. For the pouch, a shape was drawn along the apical hinge-pouch fold. For the notum, the apical hinge-notum fold and wing disc outline was used ^78^. The area between those two areas and the wing disc outline was used as hinge area. Then, the above-described ROI segmentation with measurement of Robo2-GFP intensity in ROI3 and ROI5 was applied individually for each z-section. The results were then processed as follows. Individual clones were tracked through the z-stack by their centroids and the Similarity And Distance Indices function in Past4 ^79^, resulting in up to 4 data points per clone. To avoid non genotype specific changes in Robo2-GFP intensity due to cyst formation and consequent changes of the apical plane in clones, the data points were filtered based on changes in actin levels. Data points from z-sections in which the clone had actin changes of >20% between ROI3 and ROI5, while Robo2-GFP intensity increased, were excluded. Vice versa, clones with actin changes of <-20% between ROI3 and ROI5, while Robo2-GFP intensity decreased, were excluded as well. Furthermore, small clones beneath 30µm^2^ were excluded, as well as clones which were only detected in one of the 4 z-sections. Eventually, the data point of a clone, which was used for further statistics, derived from the most apical z-section that fulfilled all those criteria.

### Illustration expression pattern

Procreate (Version 5.2.9) was used for illustration of cell surface receptor expression patterns (**Fig 1**). Characteristic points of reference, such as hinge folds, A/P and D/V boundary, were used to transfer the expression domains manually into a wing disc template. The grading of fluorescence intensity is based on overall impression of intensity across the wing disc and is presented as a subjective approximation. Bitmap tracing was done in Inkscape (0.92.4) to convert the areas of expression into vector graphics. For the overlay in **Fig S1C**, intensity grading of individual expression patterns was disregarded, and the entire expression domain was assigned the value 1 through the FIJI-threshold function. All illustrated receptors were then added together using FIJI-image calculator function. The final image was colored using LUT_Cyan-Hot.

## Author Contributions

Conceptualization FF, AF, AKC

Data curation FF, LE, AF, KH

Formal analysis FF, AF, LE, KH, DP, VW

Funding acquisition AKC

Investigation FF, AF, LE, KH, DP, VW, RB

Methodology FF, AF, LE, KH, DP, VW, RB

Project administration AKC

Supervision AKC

Validation FF, AF, LE, KH, DP, VW, RB

Visualization FF, LE

Writing original draft FF, AKC

Writing review & editing LE, AF, FF, AKC

## Acknowledgements

We thank the staff of the Life Imaging Center (LIC) in the Hilde Mangold House (HMH) of the Albert-Ludwigs-University of Freiburg for help with their confocal microscopy resources, and the excellent support in image recording. We specifically thank the DFG for supporting our imaging work through project number 414136422. We thank David Bilder, Ishwar Hariharan, George Pyrowolakis, Martin Juenger, Suzanne Eaton, Dirk Bohmann, Mirka Uhlirova, Helena Richardson, Jennifer Zallen, Ruth Johnson, Frank Schnorrer, David Strutt, Andrea Brand, Yoshimasa Yagi, Karl-Friederich Fischbach, Mel Feany for sharing reagents. We thank the Bloomington Drosophila Stock Centre (BDSC), the Vienna Drosophila Stock Collection (VDRC) and the Developmental Studies Hybridoma Bank (DSHB) and the Monoclonal Antibody Core Facility at the Helmholtz Zentrum Munich for providing fly stocks and antibodies. We also thank the SGBM and IMPRS-IE graduate schools for supporting our students.

---

**Supplementary Figure S1.**
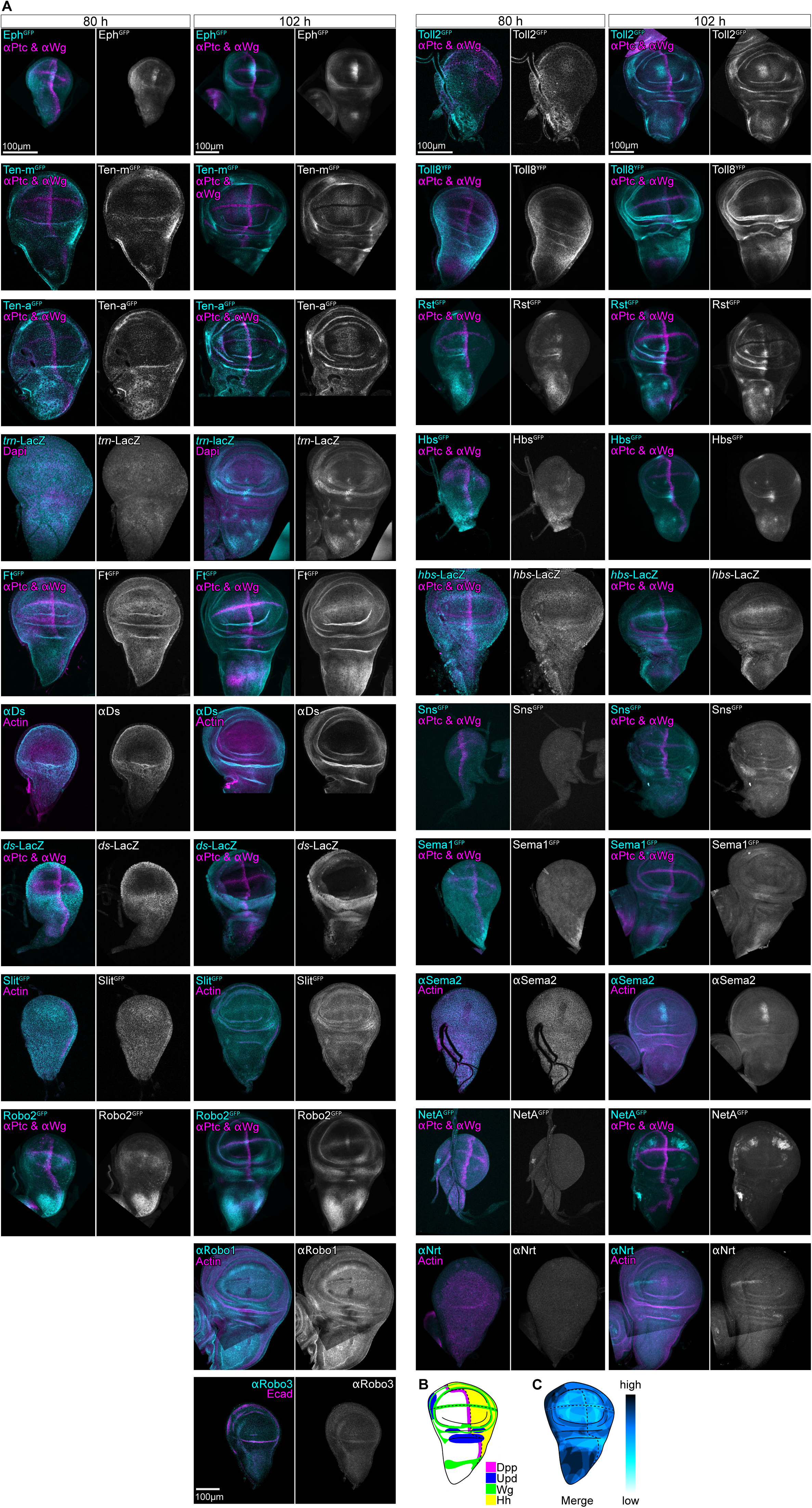
Expression patterns of cell surface molecules. **A** Wing discs expressing GFP fusion proteins and LacZ-transcriptional reporters or stained with antibodies to detect protein expression patterns at 80 h and 102 h AEL. Ptc and Wg demarcate A/P and D/V compartment boundaries. F-Actin visualizes wing disc area. All 80 h AEL discs are displayed at the same scale. 102 h discs are in a different scale. **B** Illustration of expression domains of fate specifying signaling ligands, i.e., the morphogens Dpp (magenta), Wg (green), Hh (yellow) and Upd (blue). Please note that fate specifying signaling ligands also act non-autonomously by establishing morphogen gradients, and thereby specify fate patterns in the wing discs. **C** Overlay of the illustrations of surface receptor expression pattern from (Fig 1A). The number of different receptors which are expressed is depicted by color intensity from low to high (cyan to blue).

**Supplementary Figure S2.1.**
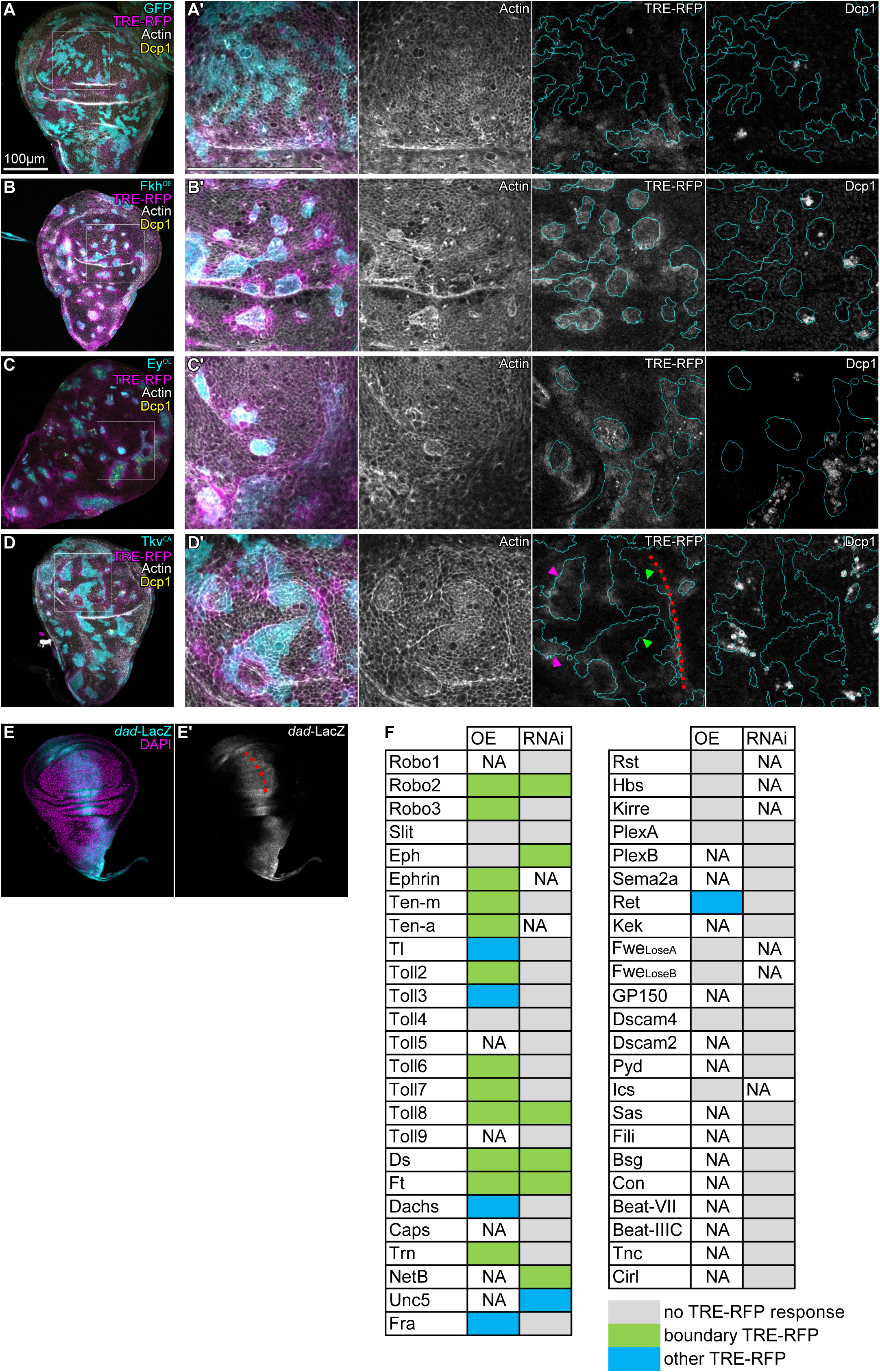
Hallmarks of interface surveillance in response to clones with aberrant cell fate. **A-D** Mosaic wing disc with wild type clones expressing *UAS-GFP* (GFP) (A), or with clonal alterations in fate specifying pathways by overexpression of Fkh (Fkh^OE^) (B), Ey (Ey^OE^) (C), or constitutively active Tkv (Tkv^CA^) (D). TRE-RFP expression is reporting JNK pathway activity (grey or magenta in A-D). Antibody staining against the cleaved effector caspase (cDcp1) visualizes apoptosis (grey or yellow in A-D). Cortical F-Actin was visualized by Phalloidin (grey). Please note that JNK activity is physiologically high at the A/P boundary in wing discs at this developmental stage. White frame marks regions shown in (A’-D’). **E** Wing disc expressing the transcriptional reporter *dad-*LacZ, a target gene of the morphogen Dpp. Red dotted line represents the A/P compartment boundary. **F** Summary of the candidate screen for JNK interface signaling induced by overexpression (^OE^) by *UAS-gene* constructs and downregulation by *UAS-gene-RNAi* (^RNAi^) constructs (see Table S2 for detailed constructs). The TRE-RFP reporter was used to access induction of JNK interface signaling at clone boundaries. Phenotypes of TRE-RFP responses were categorized in three groups: JNK interface signaling at clone boundaries (See Fig 2 for representative images), other TRE-RFP responses (for example, intra-clonal activation) and no TRE-RFP responses (phenotypes not shown). Images in A-E and A’-D’ are shown at same scale with a scale bar of 100µm.

**Supplementary Figure S2.2.**
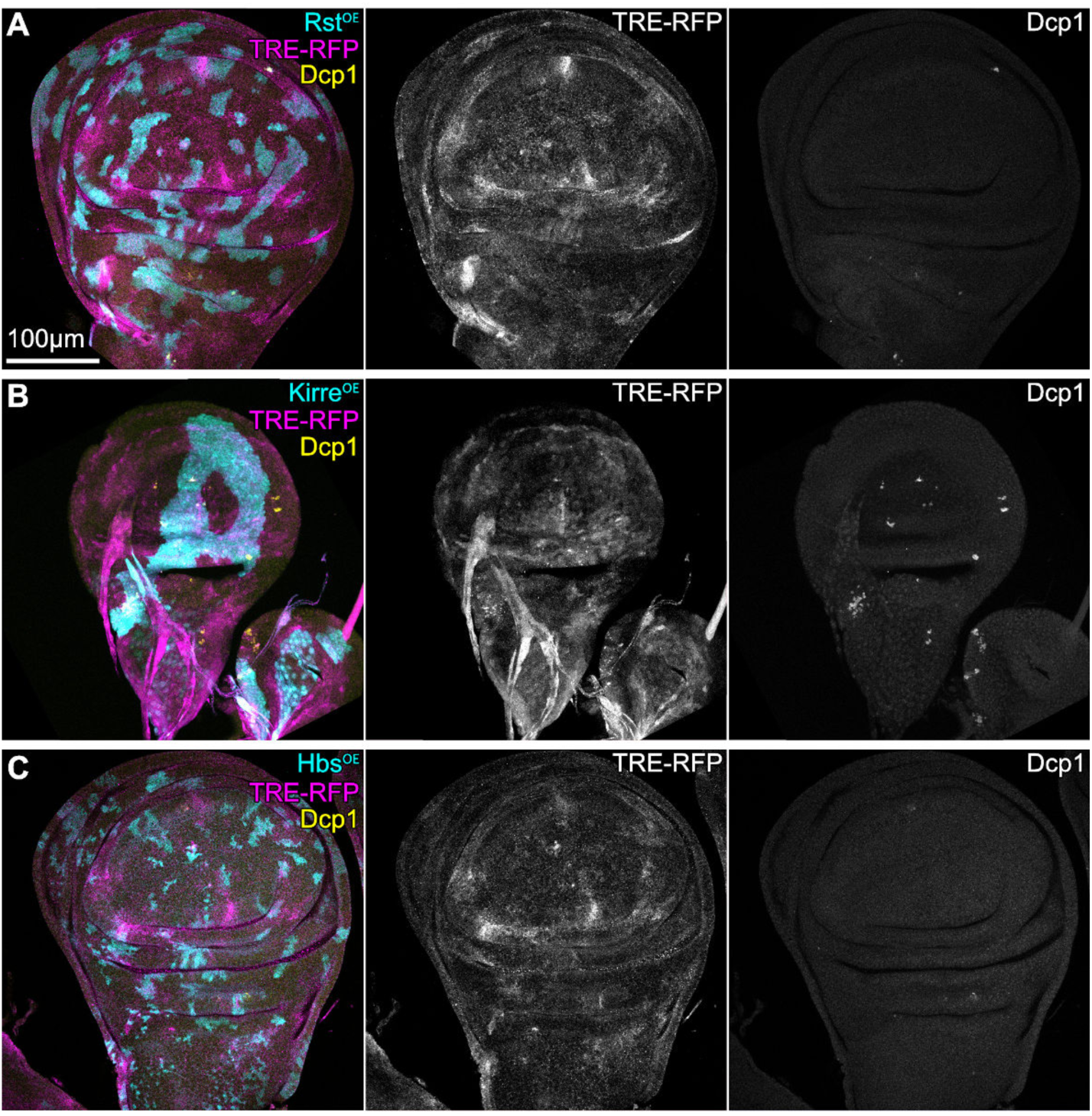
Irre cell recognition module {IRM) proteins do not induce JNK interface signaling. **A-C** Mosaic wing discs with clonal overexpression (cyan) of Rst (Rst^OE^) (A), Kirre (Kirre^OE^) (B) or Hbs (Hbs^OE^) (C) using *UAS-*constructs and analyzed for JNK activation (TRE-RFP, grey or magenta) and apoptosis (cDcp1, grey or yellow). Please note that, while Rst and Kirre appear to have a smooth clone shape, IRMs do not induce interfacial JNK activity. Images are shown at same scale.

**Supplementary Figure S3.**
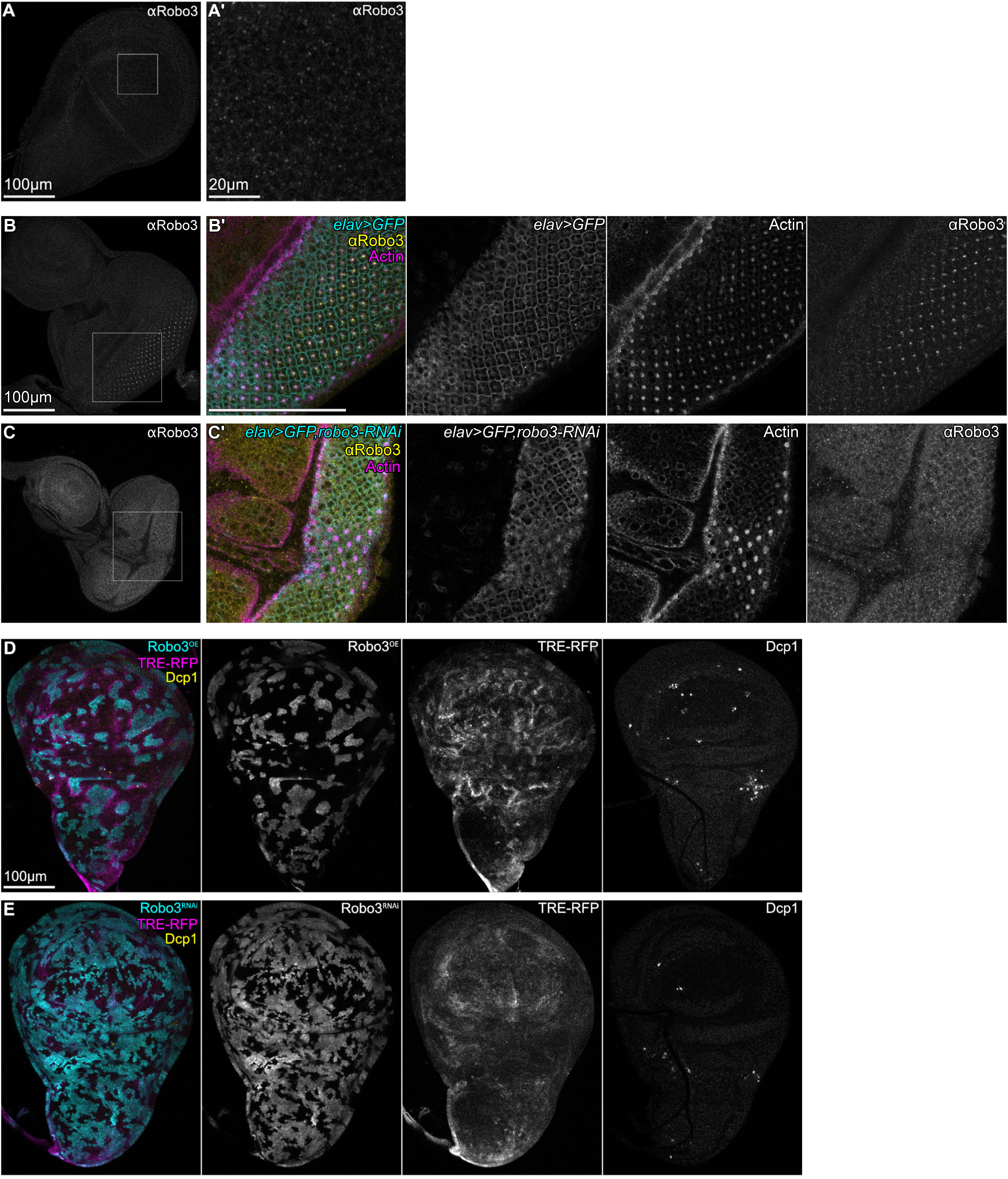
A mismatch of Robo3 receptor expression levels between neighboring cells is detected by interface surveillance in wing disc epithelia. **A** Wing imaginal disc stained by an anti-Robo3 antibody to visualize low level, punctate localization of Robo3. White frame marks region shown in (A’). **B, C** Eye imaginal discs with *elav*-GAL4 driven expression of *UAS-GFP* (E) or *UAS-GFP* and *UAS-robo3-RNAi* (F). Robo3 was visualized using an anti-Robo3 antibody. Phalloidin visualizes cortical F-Actin. Please note that Robo3 localizes to the center of assembling ommatidia clusters, which is reduced by expression of *robo3-RNAi*. White frame marks region shown in (E’, F’). Image sets are shown at the same scale in (B,C) and (B’,C’) with a scale bar of 100µm. **D, E** Mosaic wing discs where clones (cyan) either overexpress *UAS-robo3* (Robo3^OE^) (A), or express *UAS-robo3-RNAi* (Robo3^RNAi^) (B) to reduce Robo3 function. TRE-RFP expression is reporting JNK pathway activity (grey or magenta). Antibody staining against the cleaved effector caspase (cDcp1) visualizes apoptosis (grey or yellow). Images display the entire wing disc and are shown at the same scale.

**Supplementary Figure S4.**
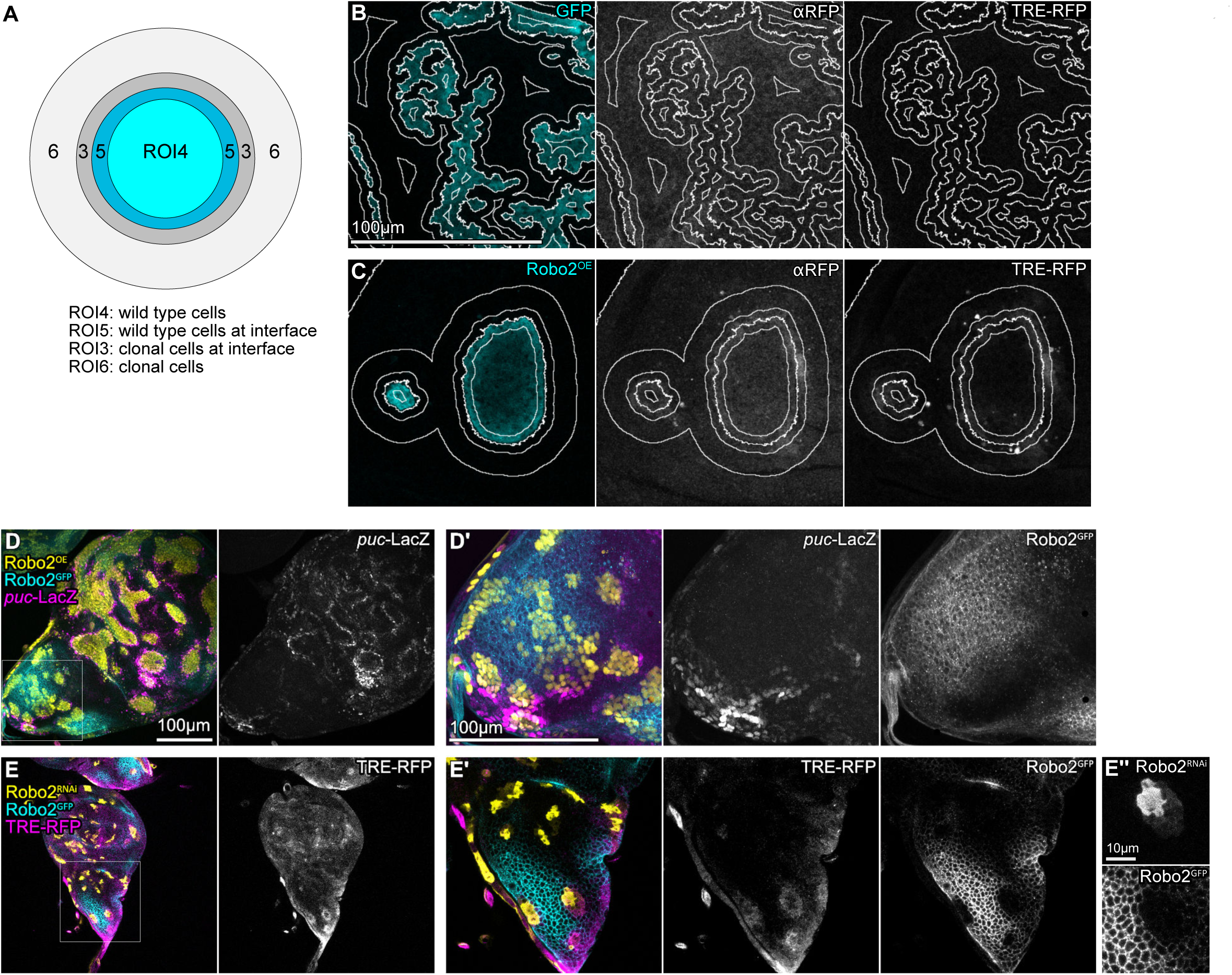
Pattern-specific interface surveillance responses. **A** Illustration of segmented regions of interests (ROIs) as also described in Fig 4F: clonal cells, that are not in touch with the interface (ROI4), a 4µm band of clonal cells (ROI5) and wild type cells (ROI3) at the interface and a 12µm band of wild type cells (ROI6) surrounding the previous ROIs (see Methods). Segmentation of the clonal interface provides the starting point for ROI annotation and subsequent measurements. **B,C** Mosaic wing discs with *UAS-GFP* expressing (D) or *UAS-robo2* (Robo2^OE^) (E) expressing clones. TRE-RFP expression is reporting JNK pathway activity. In addition to native TRE-RFP signal, anti-RFP antibody (αRFP) was used to enhance the TRE-RFP signal. White lines represent ROI segmentation. **D,E** Wing discs expressing the *robo2-GFP* fusion protein (Robo2^GFP^) under native regulatory control (grey or cyan). Mosaic clones (yellow) were induced that either overexpress *UAS-robo2* (Robo2^OE^) (D) or downregulate Robo2 using expression of *UAS-robo2-RNAi* (Robo2^RNAi^) (E). Please note the reduced Robo2 expression level within clones in (E’’). *puc*-LacZ (D) or TRE-RFP (E) expression (grey or magenta) is reporting JNK pathway activity (grey or magenta). Image sets are shown at the same scale. White frame marks region shown in (D’, E’).

**Supplementary Figure S5.**
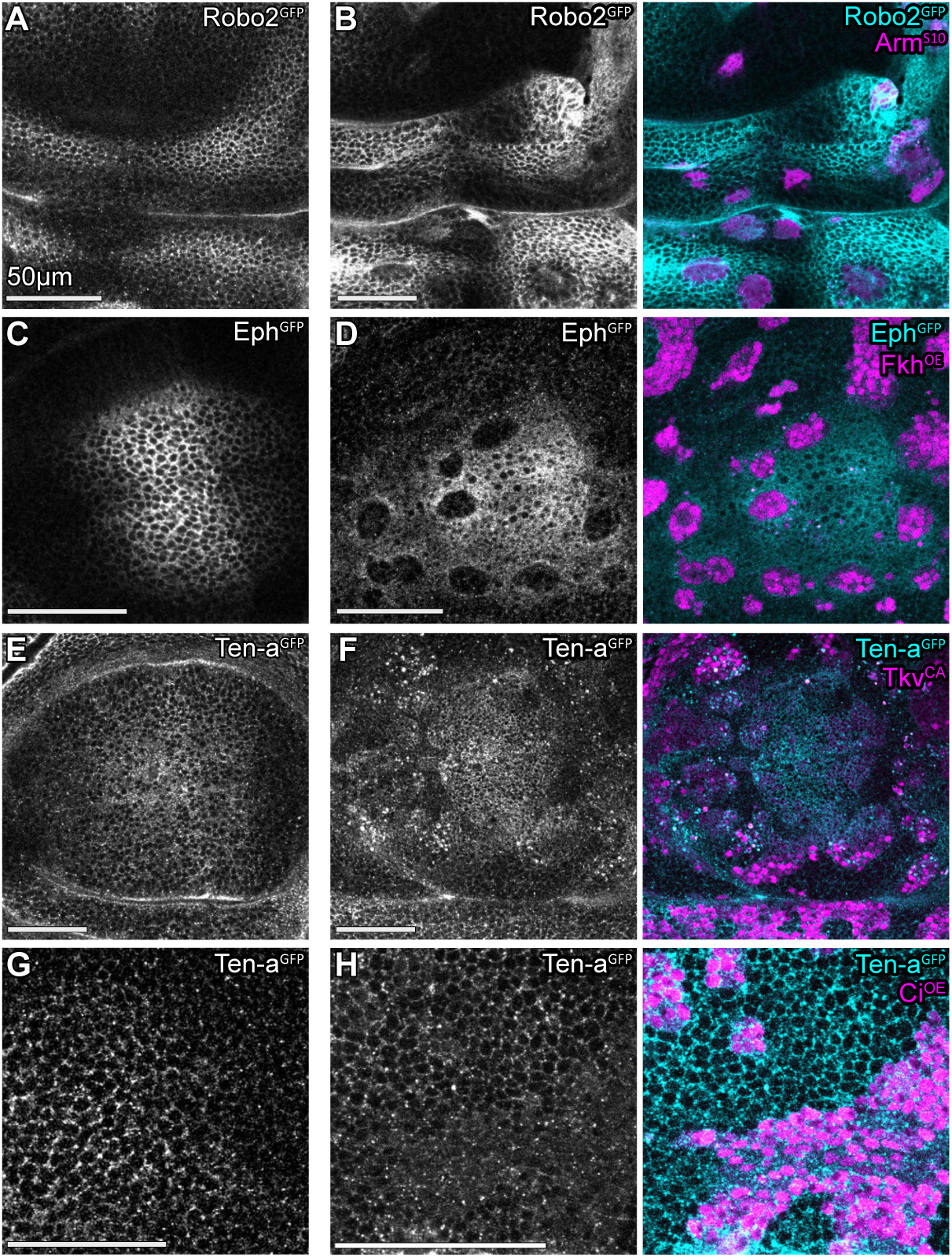
Expression of cell surface molecules in wing disc epithelia is regulated by fate-patterning pathways. **A, C, E, G** Wing discs presenting the specific expression patterns of *robo2-GFP* (Robo2^GFP^) (A), *eph-GFP* (Eph^GFP^) (C) or *ten-a-GFP* (Ten-a^GFP^) (E,G) serving as a reference for the experimentally induced changes in (B, D, F, H). **B, D, F, H** Wing discs expressing *robo2-GFP* (Robo2^GFP^) (B), *eph-GFP* (Eph^GFP^) (D) or *ten-a-GFP* (Ten-a^GFP^) (F,H) (grey or cyan). Mosaic clones (magenta) deregulate fate-specifying pathway by expression of a constitutively active Arm (Arm^S10^) (B), Fkh (Fkh^OE^) (D), constitutively active Tkv (Tkv^CA^) (F) of Ci (Ci^OE^) (H). Scale bars = 50µm.

**Supplementary Figure S6.**
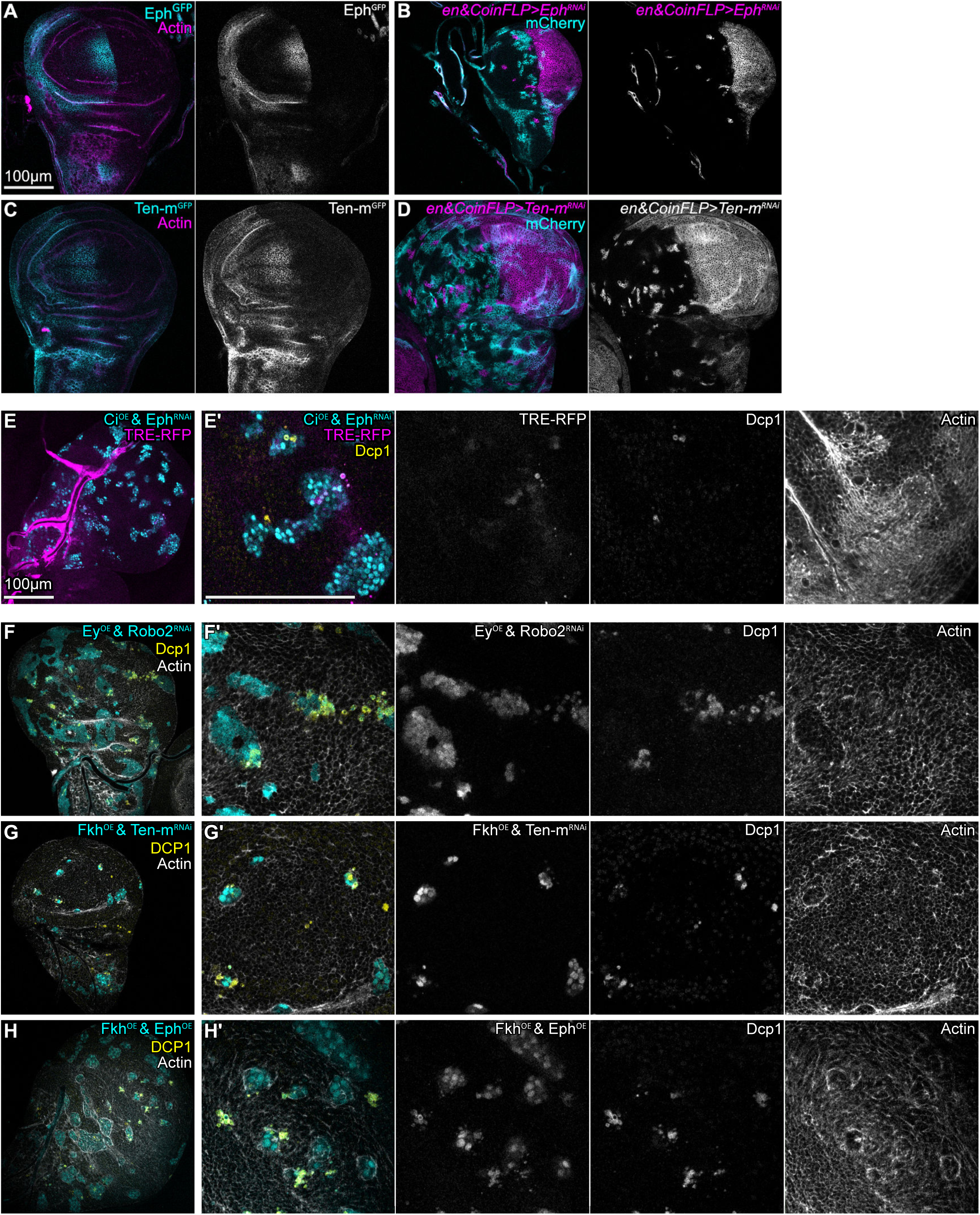
Interface surveillance is redundantly mediated by multiple cell surface receptors. **A** Wing disc from larva with the genotype *CoinFLP-LexA::GAD.GAL4/hsflp^122^ ; en-GAL4/tub>CD2>GAL4, UAS-nLac ; UAS-eph-RNAi/eph-GFP* for testing the genetic constructs used in Fig 6B. No heat shock was applied hence only *en*-GAL4 is active in the posterior compartment. Please note downregulation of Eph (grey or cyan) in the posterior compartment indicating efficient RNAi-mediated knock-down. Phalloidin visualizes cortical F-Actin (magenta). **B** Mosaic wing disc with wild type clones expressing *LexO-mCherry* (cyan) and knockdown of Eph by expression of *UAS-eph-RNAi* (Eph^RNAi^) under control of *en*-GAL4 (magenta or grey). The CoinFLP induces either LexA or GAL4 clones upon Flp-FRT recombination. *en-GAL4* drives *UAS-eph-RNAi* expression in the posterior compartment. The CoinFLP induces either LexA or GAL4 clones upon Flp-FRT recombination. Therefore GAL4/UAS clones expressing *UAS-GFP* (GFP) can also occasionally be observed in the anterior compartment (magenta). See Fig 6B. Image shown at the same scale as in (A). **C** Wing disc from larva with the genotype *CoinFLP-LexA::GAD.GAL4/+ ; en-GAL4/+ ; UAS-ten-m-RNAi/ten-m-GFP* for testing the genetic constructs used in Fig 6B. No heat shock was applied hence only *en*-GAL4 is active in the posterior compartment. Please note downregulation of Ten-m (grey or cyan) in the posterior compartment indicating efficient RNAi-mediated knock-down. Phalloidin visualizes cortical F-Actin (magenta). Image shown at the same scale as in (A). **D** Mosaic wing disc with wild type clones expressing *LexO-mCherry* (cyan) and knockdown of Ten-m by expression of *UAS-Ten-m-RNAi* (*Ten-m*^RNAi^) under control of *en*-GAL4 (magenta or grey). The CoinFLP induces either LexA or GAL4 clones upon Flp-FRT recombination. *en-GAL4* drives *UAS-Ten-m-RNAi* expression in the posterior compartment. The CoinFLP induces either LexA or GAL4 clones upon Flp-FRT recombination. Therefore GAL4/UAS clones expressing *UAS-GFP* (GFP) can also occasionally be observed in the anterior compartment (magenta). See Fig 6C. Image shown at the same scale as in (A). **E-H** Mosaic wing discs with clones deregulating a fate-specifying pathway and a cell surface receptor. Knock-down and overexpression constructs for cell surface receptors were selected to restore expression levels of the receptor within aberrant cells to wild type levels. Mosaic clones deregulating a fate-specifying pathway either express *UAS-ci* (Ci^OE^) (E), *UAS-ey* (Ey^OE^) (F) or *UAS-fkh* (Fkh^OE^) (G,H). To manipulate expression levels of cell surface receptors, we used *UAS-eph-RNAi* (Eph^RNAi^) (E), *UAS-robo2-RNAi* (Robo2^RNAi^) (F), *UAS-ten-m-RNAi* (Ten-m^RNAi^) (G) or *UAS-Eph* (Eph^OE^) (H). Please note that clones still acquire a round smooth shape. TRE-RFP expression is reporting JNK pathway activity (grey or magenta). Antibody staining against the cleaved effector caspase (cDcp1) visualizes apoptosis (grey or yellow). Phalloidin visualizes cortical F-Actin (grey). Basal sections are shown. **E’-H’** Magnified views of the pouch region of the wing disc. Images are shown at the same scale with a scale bar of 100µm.

**Supplementary Figure S7.**
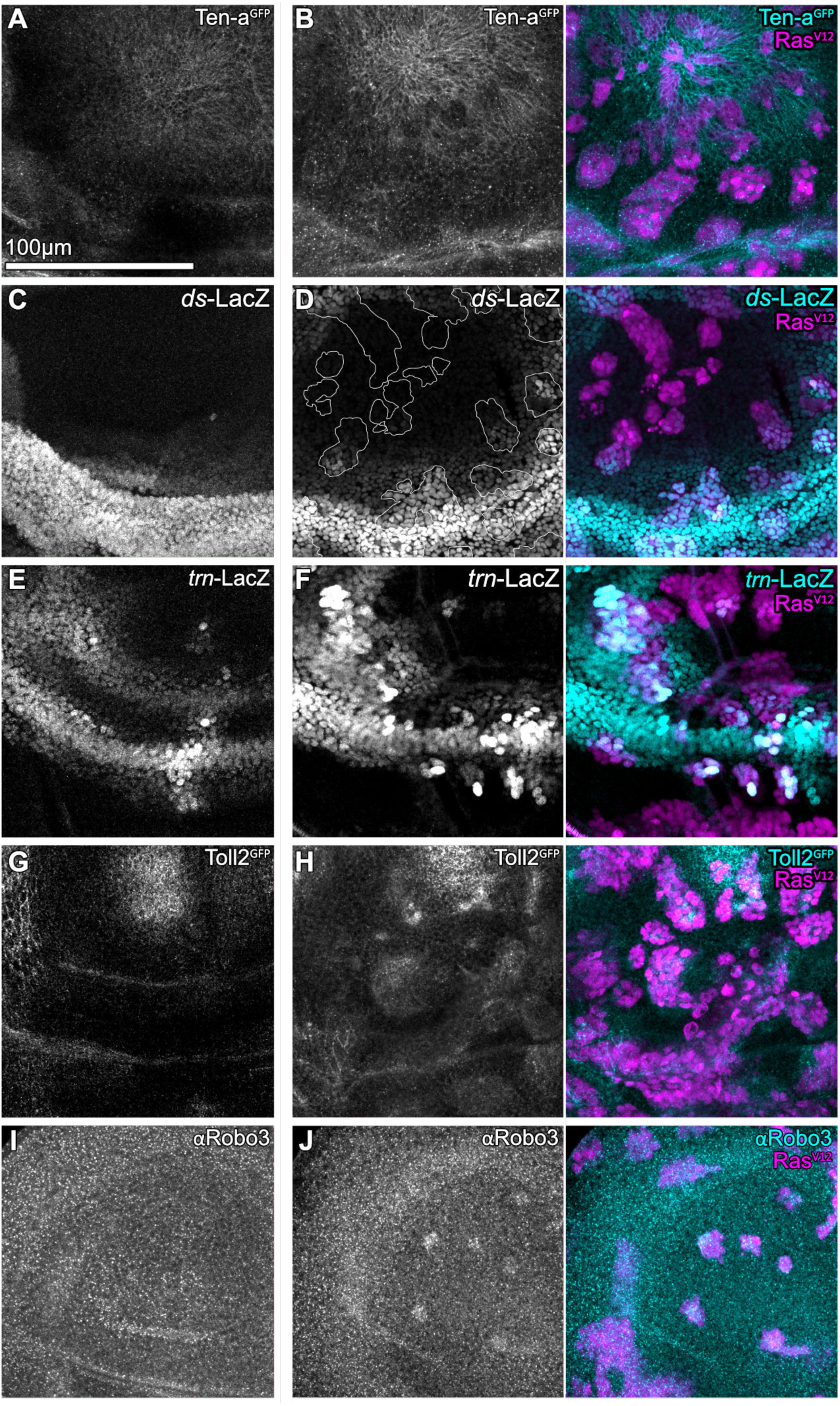
Oncogenic Ras alters cell surface molecules. **A,C,E,G,I** Wing discs presenting the specific expression patterns of *ten-a-GFP* (Ten-a^GFP^) (A)*, ds-LacZ (C), trn-LacZ (E), toll-2-GFP* (Toll2^GFP^) (G) and Robo3 (αRobo3) (I) serving as a reference for the experimentally induced changes in (B,D,F,H,J). **B,D,F,H,J** Wing discs expressing *ten-a-GFP* (Ten-a^GFP^) (B)*, ds-LacZ* (D), *trn-LacZ* (F), *toll-2-GFP* (Toll2^GFP^) (H) or being stained for Robo3 (αRobo3) (I) (grey or cyan) and carrying mosaic clones (magenta) expressing *UAS-RasV12* (Ras^V12^). Images are shown at the same scale.

**Supplementary Figure S8.**
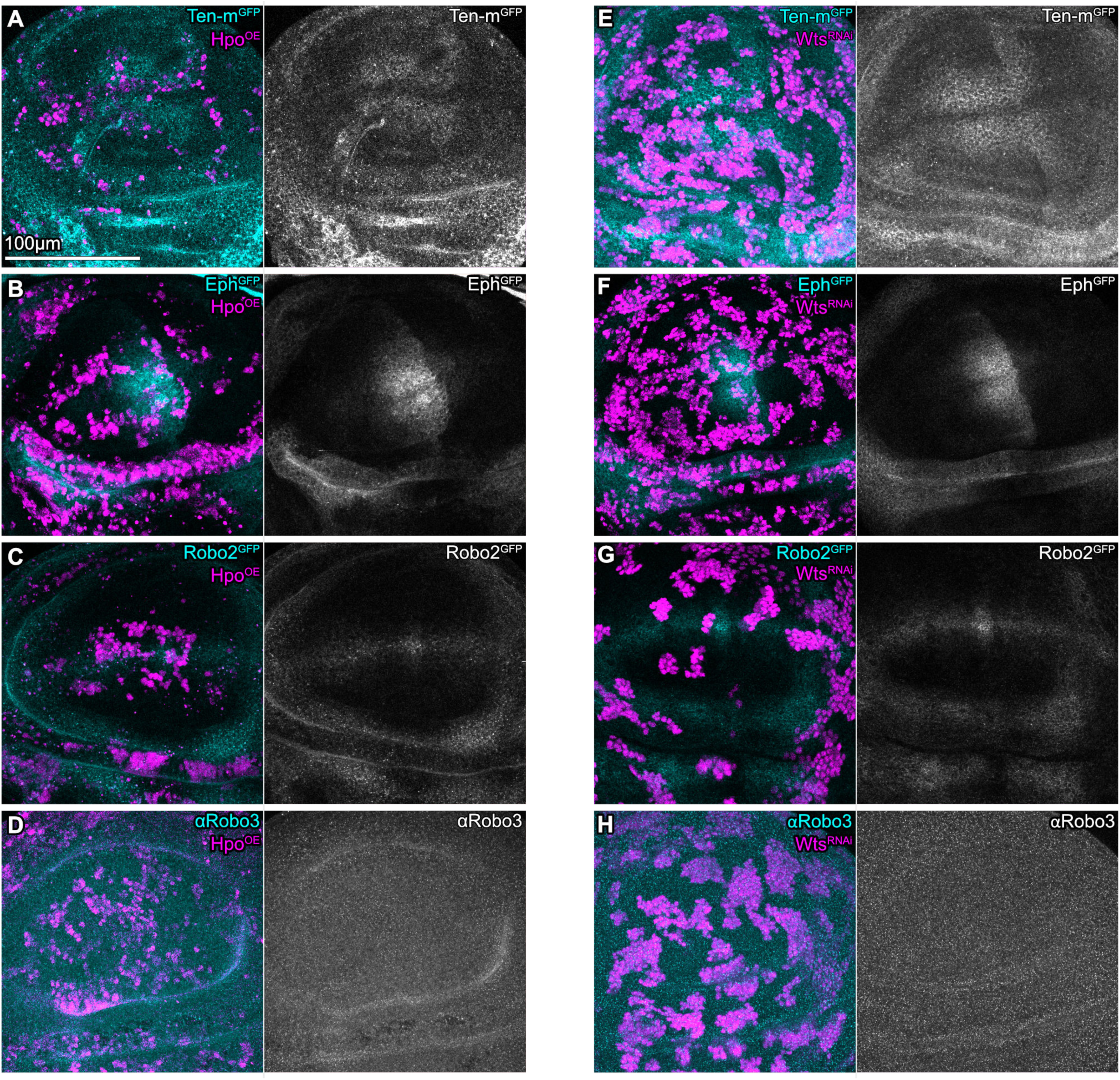
Classical models of cell-cell competition do not regulate the cell surface code. **A-H** Wing discs expressing *ten-m-GFP* (Ten-m^GFP^) (A), *eph-GFP* (Eph^GFP^) (B)*, robo2-GFP* (Robo2^GFP^) (C), stained for Robo3 (D) and carrying mosaic clones which overexpress Hippo (Hpo^OE^). Hippo overexpression reduces Yorkie function and thereby gives rise to loser cells. **A-H** Wing discs expressing *ten-m-GFP* (Ten-m^GFP^) (E), *eph-GFP* (Eph^GFP^) (F)*, robo2-GFP* (Robo2^GFP^) (G), stained for Robo3 (H) and carrying mosaic clones which express *UAS-wts-RNAi* (*wts^RNAi^)*. Knocking down *warts* function gives rise to supercompetitor/winner cells.

**Supplementary Figure S9.1.**
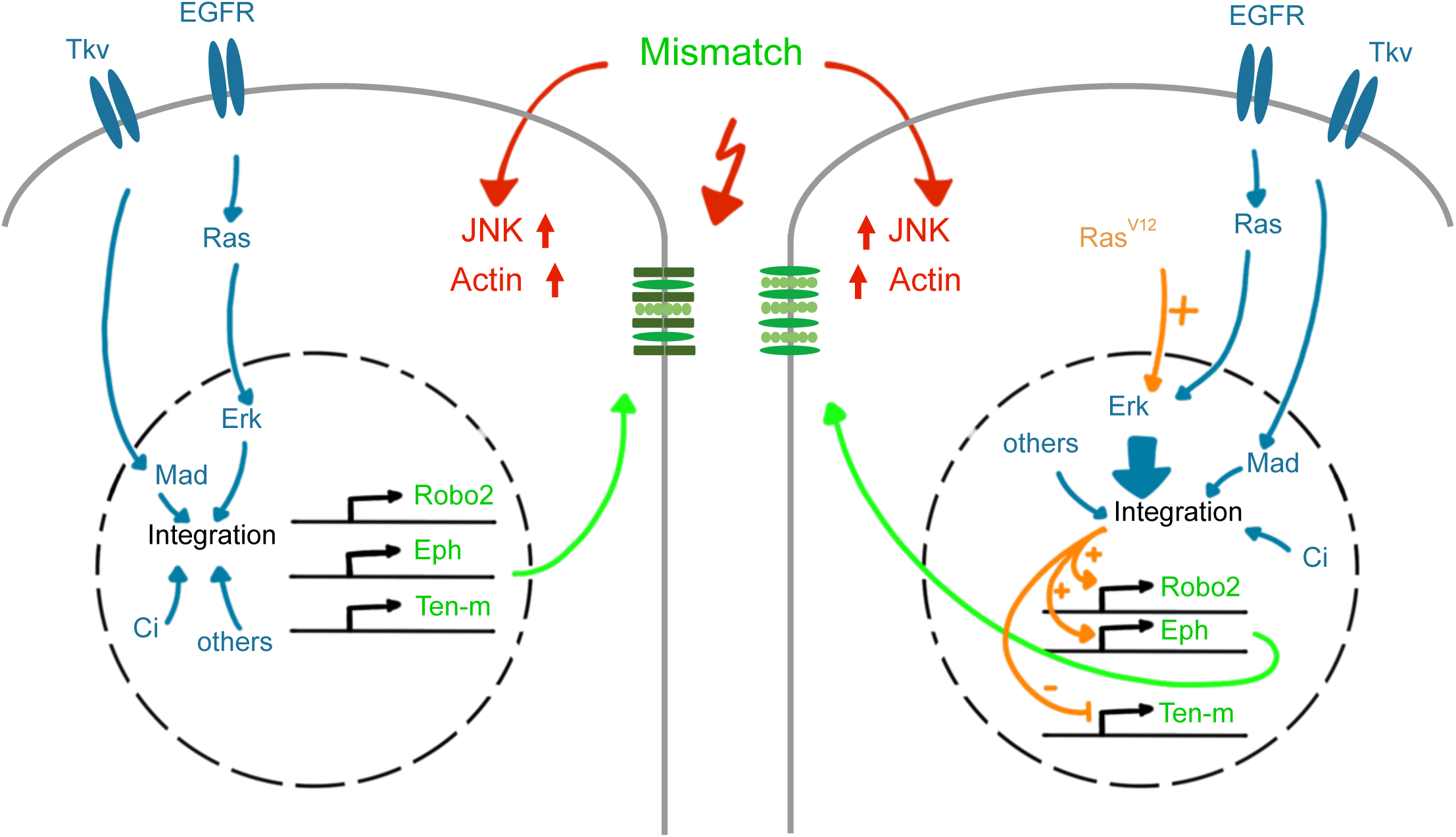
Cell surface molecules convey interface surveillance. Signaling by different developmental patterning pathways (blue) is integrated in a cell-type specific manner to drive expression of a set of genes encoding for surface molecules, which are known to play a role in neuronal guidance and targeting (green). Thus, the composition of this set of molecules displayed at the cell surface depends on the signaling environment and fate specification pathways at any specific spatial position within the imaginal disc. Aberrant cells, for example cells expressing oncogenic Ras^V12^, disrupt this position-specific regulation of gene expression (orange), leading to a changed molecular surface composition relative to that of the neighboring wild type cell. The molecular mismatch in cell surface molecules between neighboring cells induces all hallmarks of interface surveillance (red).

**Supplementary Figure S9.2.**
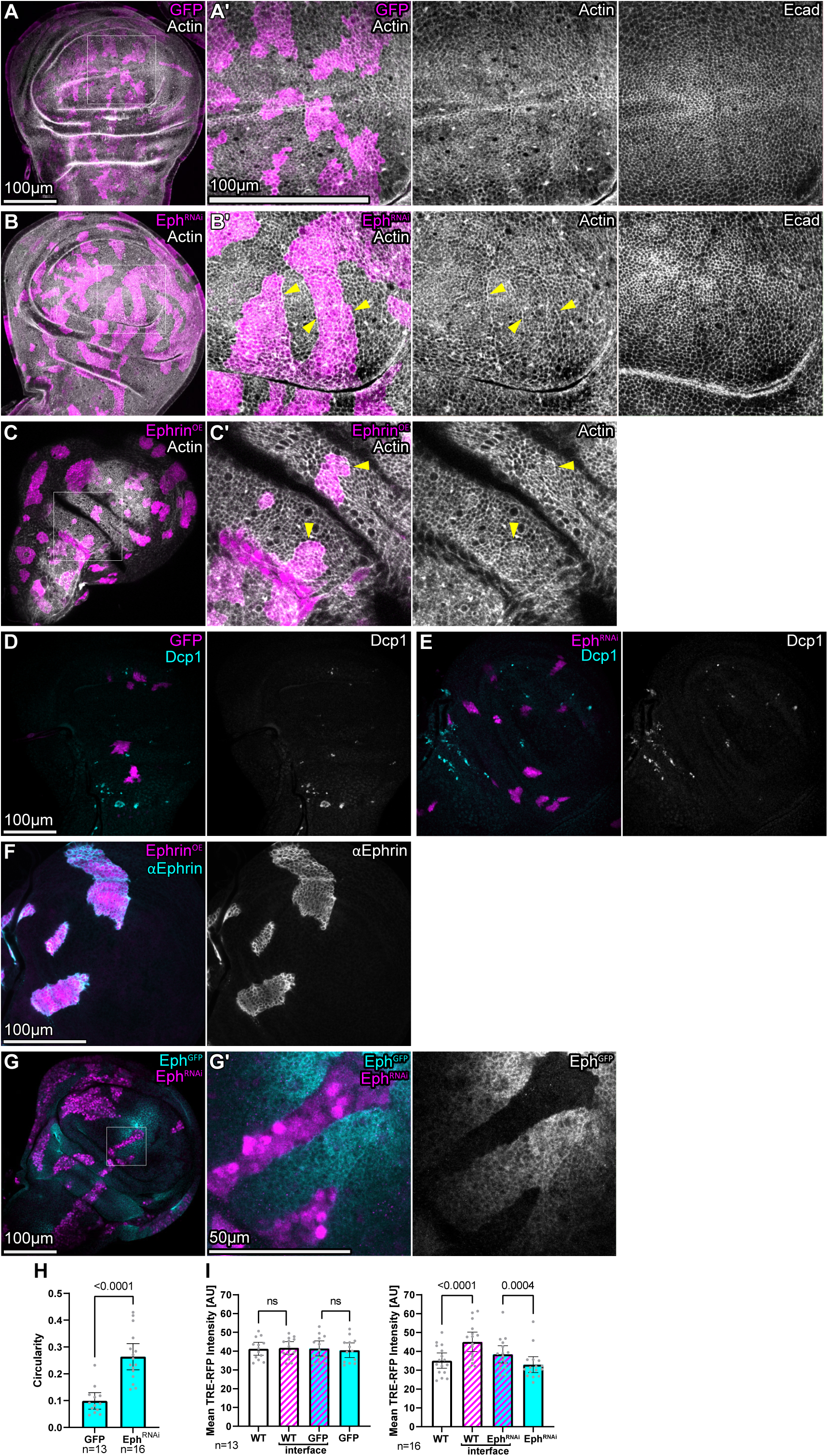
Eph receptors induce interface surveillance response. **A** A mosaic wing disc carrying wild type clones expressing *UAS-GFP* (GFP, magenta). The Fiji local-z-projector was used to project the junctional plane, as identified by E-cadherin (Ecad, grey) staining. White frame marks region shown in (A’). **B** Mosaic wing disc carrying clones expressing *UAS-eph-RNAi* (Eph^RNAi^) (magenta) to induce downregulation of Eph-receptor. The Fiji local-z-projector was used to project the junctional plane, as identified by E-cadherin (Ecad, grey) staining. Please note enrichment of F-actin at the clone interface (yellow arrows). White frame marks region shown in (B’). Image shown at the same scale as in (A). **C** Mosaic wing disc carrying clones expressing UAS-ephrin (Ephrin^OE^) to express high levels of Ephrin. An apical section is shown. Please note enrichment of F-actin at the clone interface (yellow arrows). White frame marks region shown in (C’). Image shown at the same scale as in (A). **D, E** Mosaic wing disc carrying mosaic clones expressing only *UAS-GFP* (GFP, magenta) or clones also expressing *UAS-eph-RNAi* (Eph^RNAi^) to induce downregulation of Eph-receptor. Antibody staining against the cleaved effector caspase (cDcp1) visualizes apoptosis (grey or cyan). Basal sections are shown. **F** Mosaic wing disc with clones expressing *UAS-ephrin* (Ephrin^OE^) and stained with an antibody against Ephrin (αEphrin). **G** A wing disc expressing *eph-GFP* (Eph^GFP^) and carrying mosaic clones *UAS-eph-RNAi* (Eph^RNAi^) to induce downregulation of the Eph-receptor. White frame marks region shown in (G’). Please note the strong downregulation of Eph in clones, demonstrating efficiency of knock-down using *UAS-eph-RNAi* (Eph^RNAi^). **H** Graph depicting circularity observed for individual *UAS-GFP* (GFP) and *UAS-eph-RNAi* (Eph^RNAi^) expressing clones. Sample number (n) for individual wing discs and p-values of a two tailed, unpaired t-test are displayed in graphs. **I** Graphs depicting mean fluorescence intensity of TRE-RFP reporter in zones of measurement around clones, as depicted in Fig 4E. Graphs display results for mosaic discs with *UAS-GFP* labeled clonal cells (left) or containing *UAS-eph-RNAi* (Eph^RNAi^) expressing clones (right). Sample number (n) for individual wing discs and p-values of a repeated measured one-way ANOVA with Tukey’s multiple comparisons test are displayed in graphs.

**Table S1.**
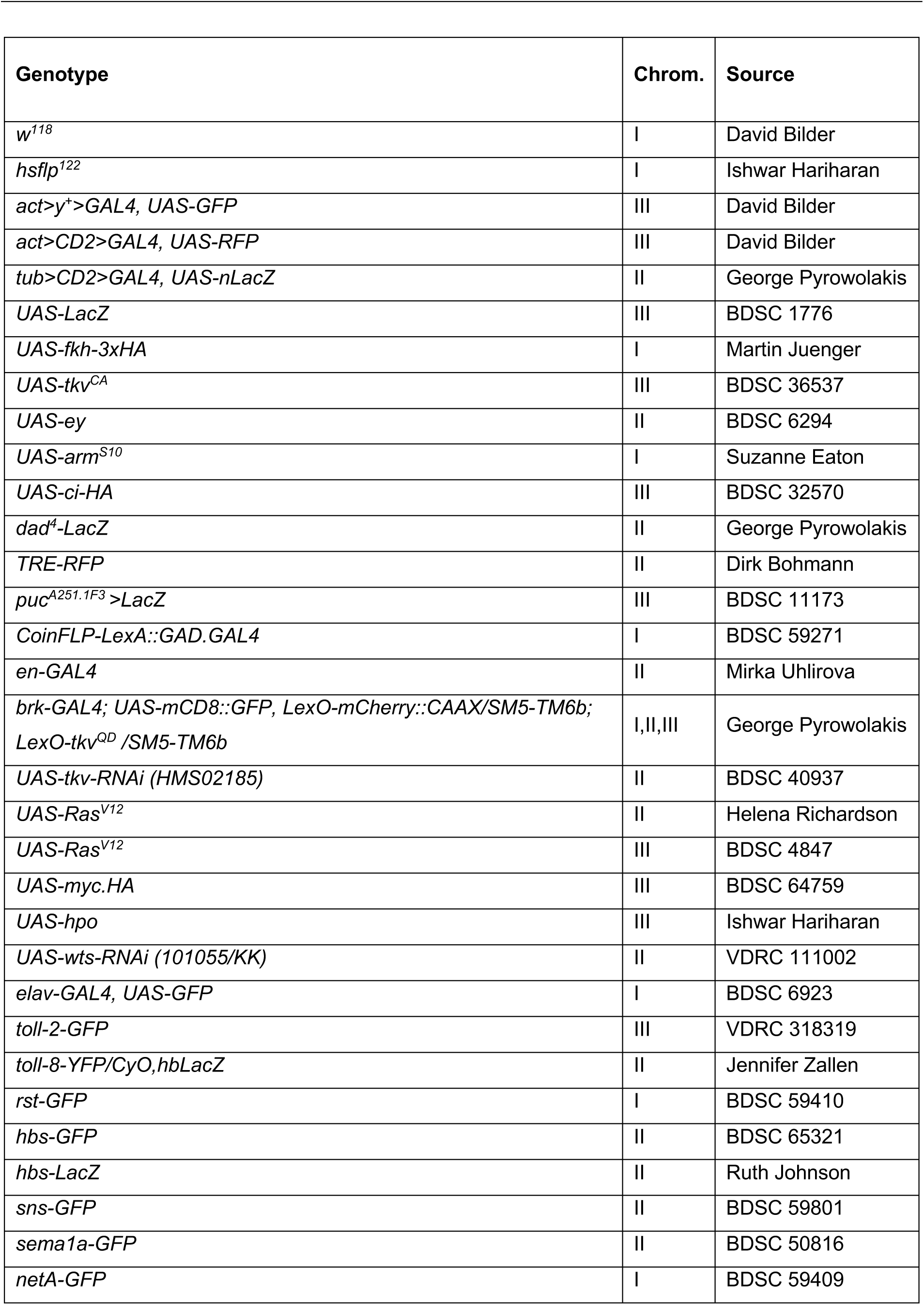
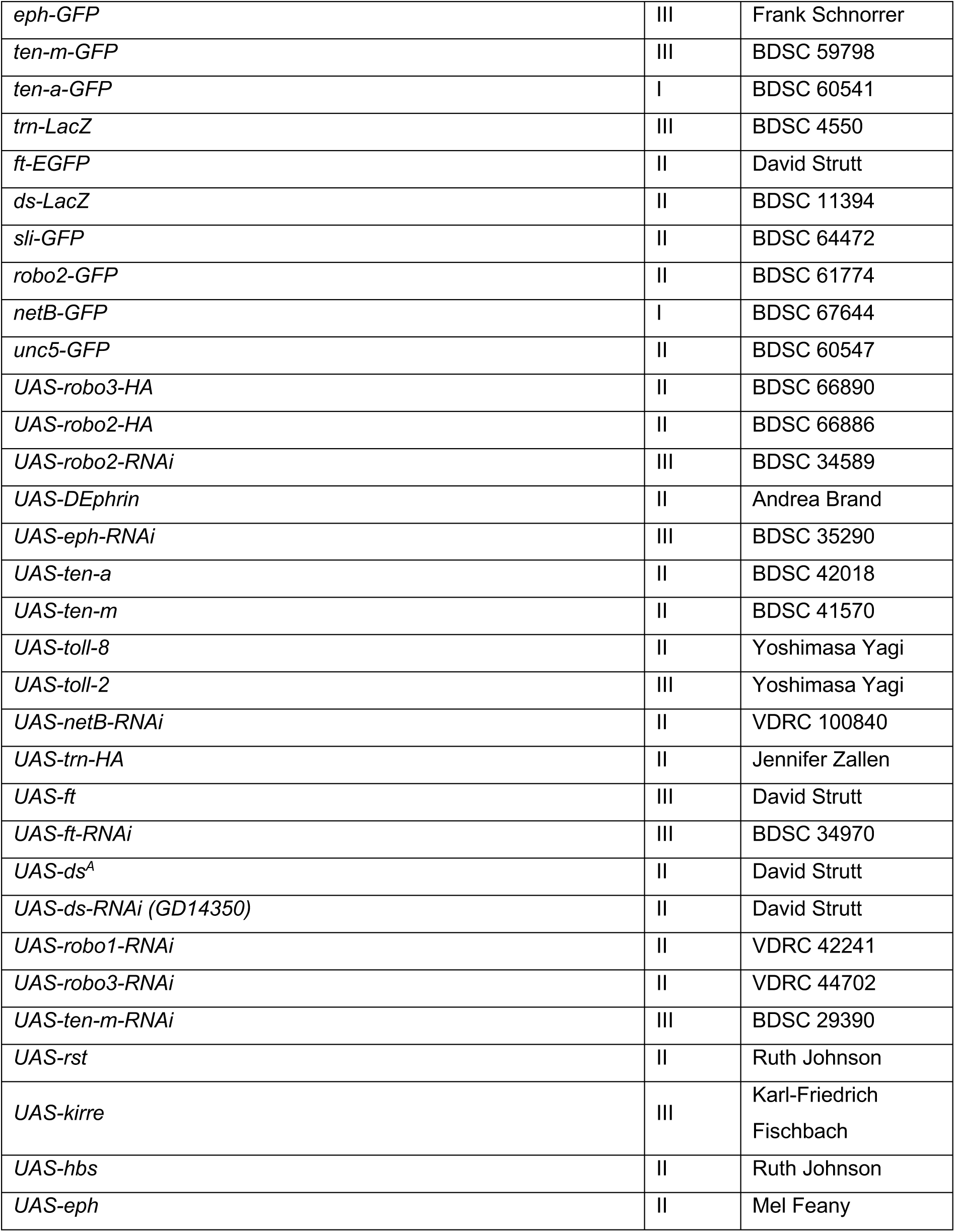
Fly strains.

**Table S2.** Additional Fly strains for candidate genetic screen for interface JNK signaling in Figure S2.

| **Genotype** | **Source** |
| --- | --- |
| *UAS-plexA-RNAi*<br>(HM05221, GD/14483, GL01870, KK/101499) | BDSC 30483, VDRC 27240,<br>BDSC 67845, VDRC 107004 |
| *UAS-plexB-RNAi* (HM05122, GD/16420) | BDSC 28911, VDRC 46687 |
| *UAS-sema2a-RNAi* (GL00356) | BDSC 35432 |
| *UAS-plexA* | BDSC 8479 |
| *UAS-sli.D* | BDSC 67689 |
| *UAS-slit-RNAi* (KK/100803) | VDRC 108853 |
| *UAS-cirl-RNAi*<br>(KK/108383, JF02674, HMS00136) | VDRC100749,<br>BDSC 27524, BDSC 34821 |
| *UAS-tl* | BDSC 58987 |
| *UAS-tl-RNAi* (JF01276, JF01491) | BDSC 31477, BDSC 31044 |
| *UAS-toll-2-RNAi* | BDSC 30498 |
| *UAS-toll-3* | Yoshimasa Yagi |
| *UAS-toll-3-RNAi* (HM05012) | BDSC 28526 |
| *UAS-toll-4* | Yoshimasa Yagi |
| *UAS-toll-4-RNAi* (HM050297) | BDSC 28543 |
| *UAS-toll-5-RNAi* (HM05212) | BDSC 29533 |
| *UAS-toll-6* | Yoshimasa Yagi |
| *UAS-toll-6-RNAi* (HMC0584, HMS04251) | BDSC 64968, BDSC 56048 |
| *UAS-toll-7* | Yoshimasa Yagi |
| *UAS-toll-7-RNAi* (HM05230) | BDSC 30488 |
| *UAS-toll-8-RNAi* | BDSC 28519 |
| *UAS-toll-9-RNAi* (HM05227, HMS00171) | BDSC 30535, BDSC 34853 |
| *UAS-dachs.V5* | BDSC 28814 |
| *UAS-dachs-RNAi* (JF02743) | BDSC 27664 |
| *UAS-caps-RNAi*<br>(GD/2530, GD/14433, JF02854) | VDRC 3046,<br>VDRC 27097, BDSC 28020 |
| *UAS-trn-RNAi*<br>(GLC01638, KK/104688, HM05011) | BDSC 50520,<br>VDRC 107883, BDSC 28525 |
| *UAS-netB-RNAi* (KK/103672) | VDRC 100840 |
| *UAS-unc5-RNAi* (HMS01099) | BDSC 33756 |
| *UAS-fra* | BDSC 8814 |
| *UAS-fra-RNAi* (JF01231, JF01457) | BDSC 31469, BDSC 31664 |
| *UAS-kek1-RNAi* (HMC04442) | BDSC 57000 |
| *UAS-kek3-RNAi* (HMS05948) | BDSC 77354 |
| *UAS-kek4-RNAi (HMC06307)* | BDSC 67206 |
| *UAS-kek5-RNAi (HMS01996)* | BDSC 40830 |
| *UAS-ret.L* | BDSC 59002 |
| *UAS-ret-RNAi (HMC04143)* | BDSC 55872 |
| *UAS-fwe<sup>LoseB</sup>* | Eduardo Moreno |
| *UAS-fwe<sup>LoseA</sup>* | Eduardo Moreno |
| *UAS-GP150 (HMS00395)* | BDSC 32400 |
| *UAS-dscam4-RNAi (HMC03277)* | BDSC 51508 |
| *UAS-dscam4* | BDSC 17117 |
| *UAS-dscam2-RNAi (HMC03411)* | BDSC 51839 |
| *UAS-pyd-RNAi* | Ruth Johnson |
| *UAS-ics.O* | BDSC 66567 |
| *UAS-sas-RNAi (KK/106177)* | VDRC 100901 |
| *UAS-fili-RNAi (HM05054)* | BDSC 28568 |
| *UAS-bsg-RNAi (HMC03195)* | BDSC 52110 |
| *UAS-con-RNAi (HM05178)* | BDSC 28967 |
| *UAS-beat-VII-RNAi (HMC05049)* | BDSC 60056 |
| *UAS-beat-IIIc-RNAi (JF03286)* | BDSC 29607 |
| *UAS-tnc-RNAi (HMC05051)* | BDSC 60058 |

**Table S3.**
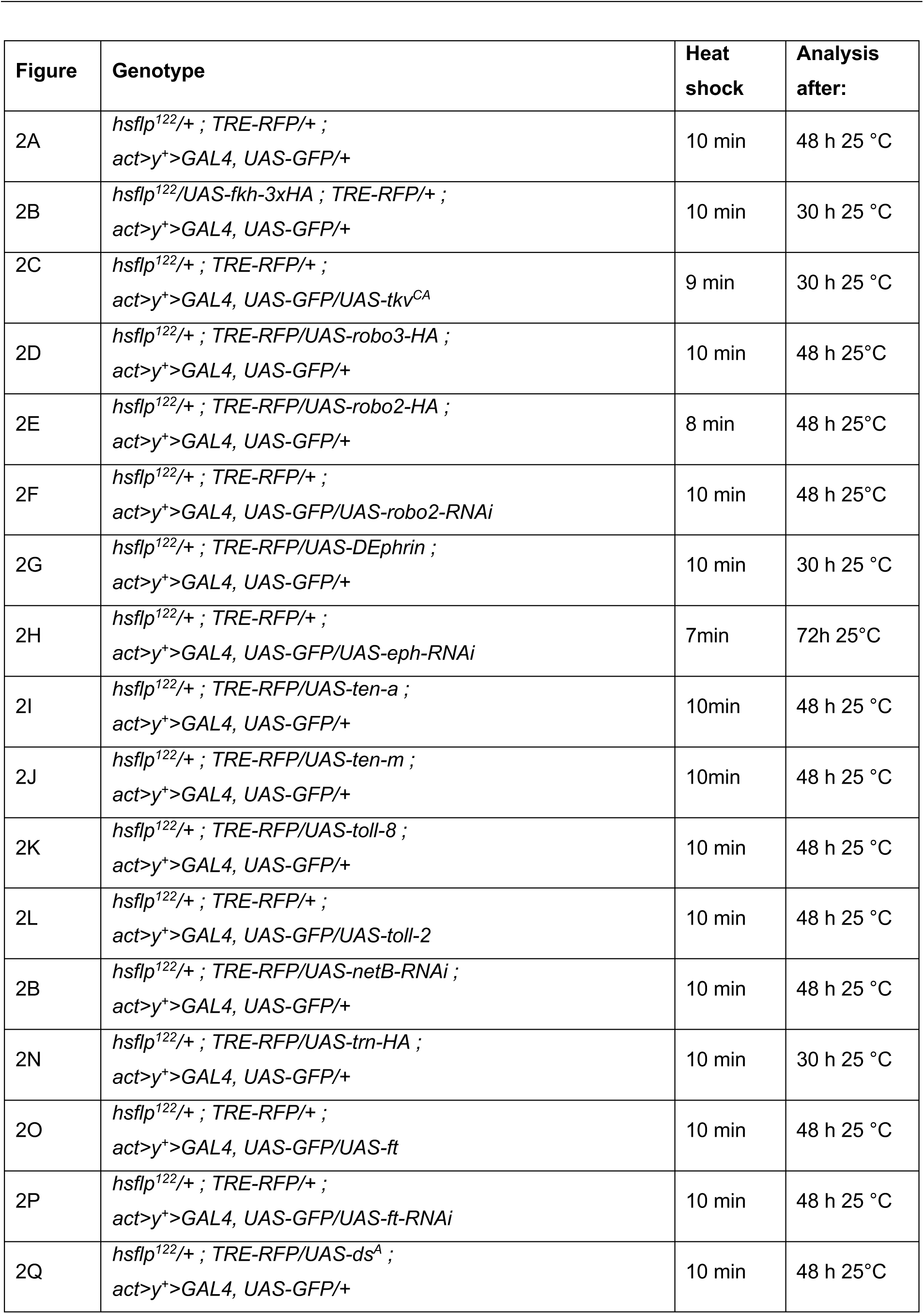
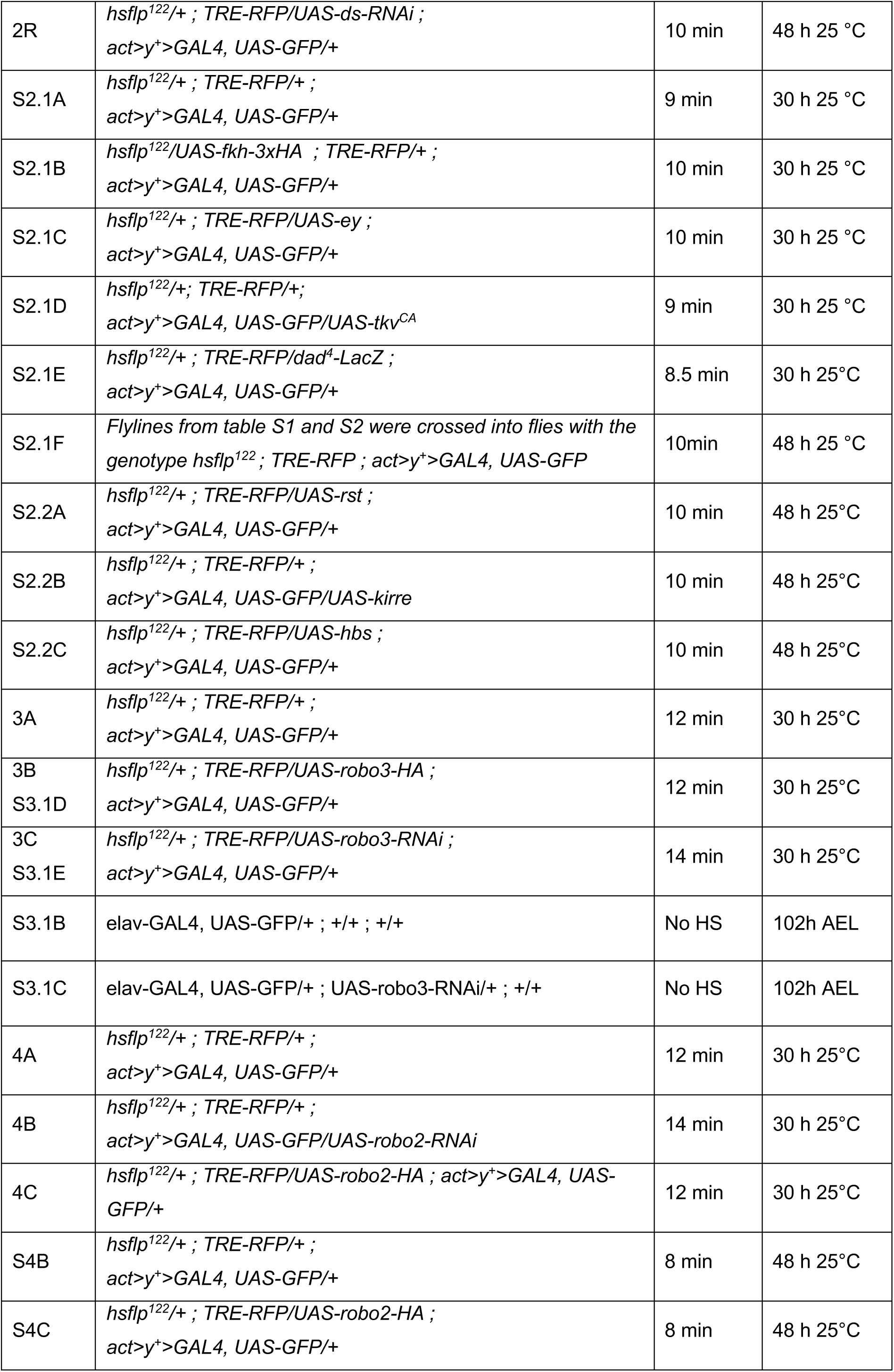
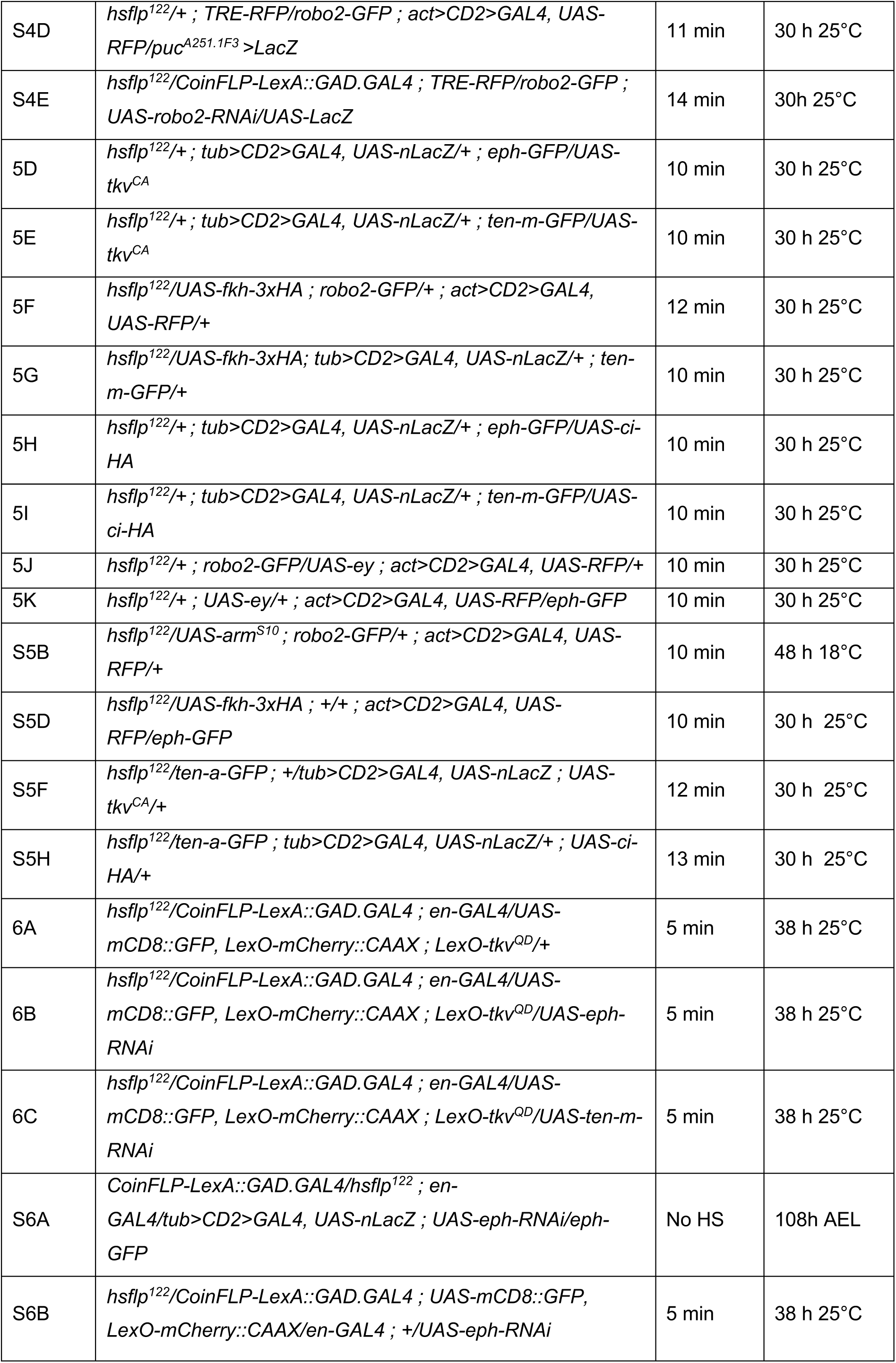
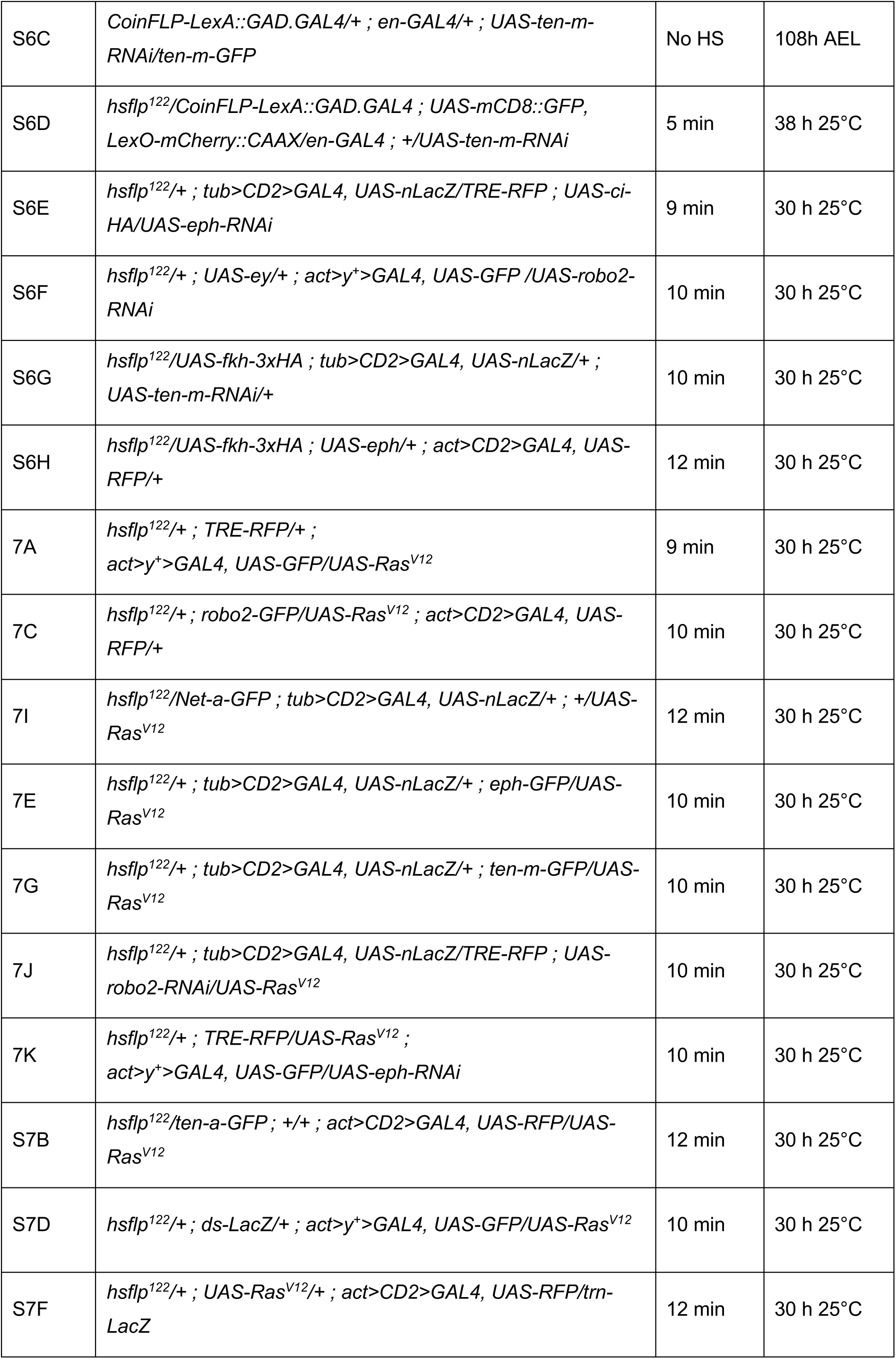
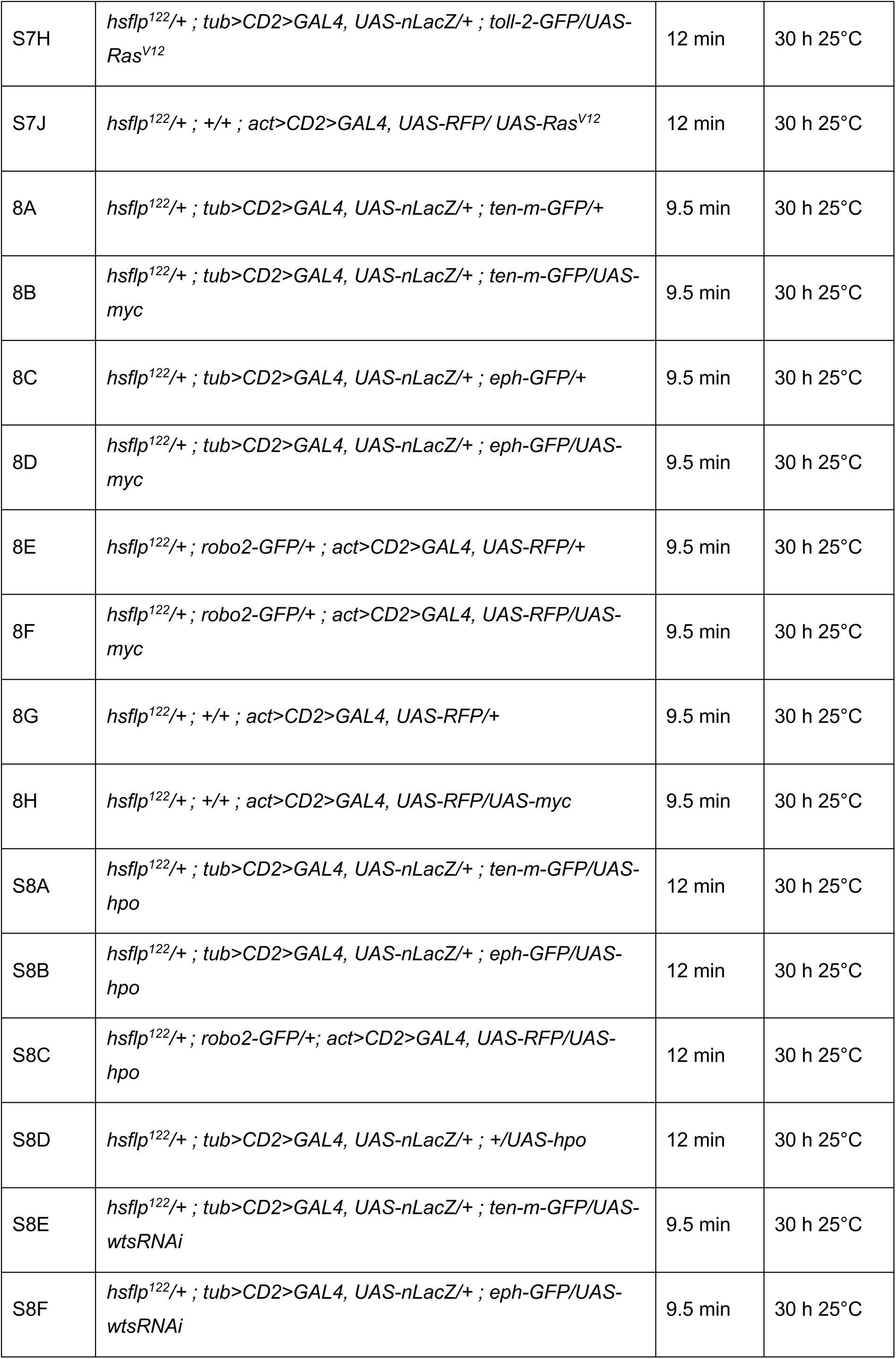
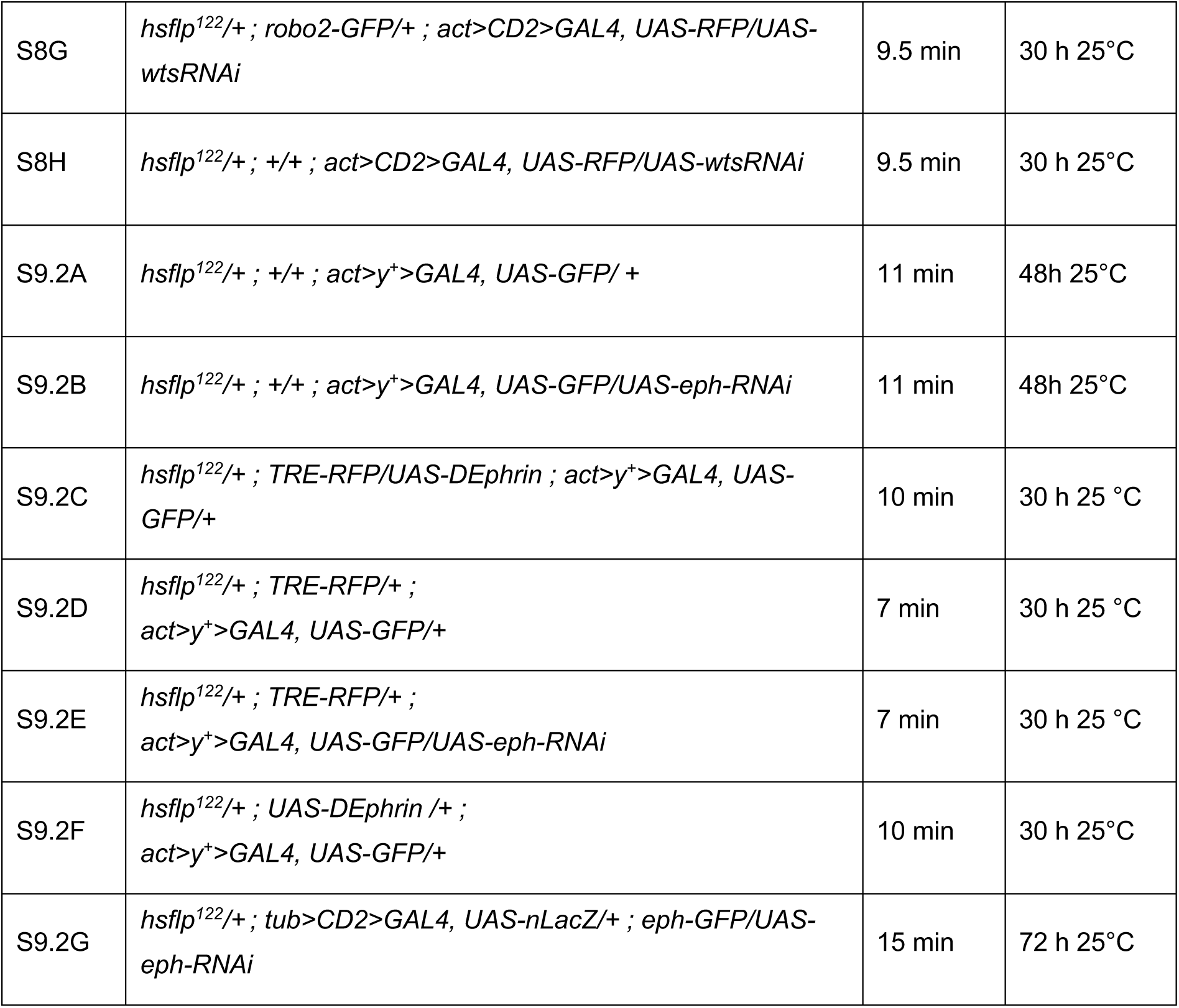
Detailed genotypes per figure panel.

**Table S4.** Selected z-positions for figure panels with overlay of channels from different z-positions. In most figure panels, individual channels represent the same z-position from a confocal image stack. However, some panels may assemble channels from different z-position in the wing disc (details below). Such a portrayal was chosen to better visualize the spatially distinct phenotypes at different z-positions within the tissue, specifically of (1) junctional actin (apical), (2) cytoplasmic/nuclear RFP (lateral) and (3) cDcp1 (basal) in one image panel. This was meant to reduce the data load in the manuscript that would be otherwise required to visualize all channels at all z-positions. All full original image stacks are of course available upon request.

|  |  |
| --- | --- |
| S2.1A' | *Overlay:* apical z-sections of actin and clonal marker. Basal z-sections of TRE-RFP.<br>*Black and white images:* apical z-section of Actin. Basal z-sections of TRE-RFP and Dcp1. |
| S2.1B' | *Overlay:* local Z projection of apical actin, apical clonal marker, and apical to lateral TRE-RFP.<br>*Black and white images:* local z projection of apical actin, basal max projection of TRE-RFP and Dcp1 |
| S2.1C' | *Overlay:* apical z-sections of actin, clonal marker and TRE-RFP.<br>*Black and white images:* apical z-section of actin and lateral z-section of TRE-RFP and Dcp1. |
| S2.1D' | *Overlay:* max projection of apical z-sections of actin, clonal marker and TRE-RFP.<br>*Black and white images:* apical z-section of actin, lateral z-section of TRE-RFP and basal z-section of Dcp1. |
| 3A and 3A' | Basal z-section. |
| 3A'' | Max projection of 3 apical z-sections. |
| 3B and 3B' | Lateral z-section of clonal marker and TRE-RFP. Basal z-section of Dcp1. |
| 3B'' | Max projection of 3 apical z-sections. |
| 3C and 3C' | Lateral z-section of clonal marker and TRE-RFP. Basal z-section of Dcp1. |
| 3D'' | Max projection of 3 apical z-sections. |
| 4A | Lateral z-section for clonal marker and TRE-RFP (different z-section for pouch and notum). Max projection of z-sections for Dcp1. |
| 4A' and 4A'' | Max projection of apical z-sections for Actin, a lateral z-section for TRE-RFP and clonal marker and max projection of z-sections for Dcp1. |
| 4B | Lateral z-section. |
| 4B' | Apical z-section for Actin and clonal marker. Lateral z-section for TRE-RFP and Dcp1. |
| 4B'' | Max projection of lateral to basal z-sections. |
| 4C | Basal z-section for TRE-RFP and clonal marker (different z-section for pouch and notum). Max projection of all z-sections for Dcp1. |
| 4C' | max projection of 2 apical z-sections for Actin, TRE-RFP and clonal marker. Basal z-section for Dcp1. |
| 4C'' | max projection of 2 apical z-sections for Actin. Lateral z-section for clonal marker and TRE-RFP. Max projection of all z-sections for Dcp1. |
| 6A-C' | lateral z-section for LexA and Gal4 clonal marker and Actin. Max projection of basal z-sections for Dcp1. |
| S6E' | Basal z-section in clonal marker, TRE-RFP, Dcp1. Apical z-section for Actin. |
| 7H and 7I | Max projection of apical to lateral z-section in the hinge |
| 8 and S8 | Local z projection of lateral z-section |

